# Comprehensive mapping of tissue cell architecture via integrated single cell and spatial transcriptomics

**DOI:** 10.1101/2020.11.15.378125

**Authors:** Vitalii Kleshchevnikov, Artem Shmatko, Emma Dann, Alexander Aivazidis, Hamish W King, Tong Li, Artem Lomakin, Veronika Kedlian, Mika Sarkin Jain, Jun Sung Park, Lauma Ramona, Elizabeth Tuck, Anna Arutyunyan, Roser Vento-Tormo, Moritz Gerstung, Louisa James, Oliver Stegle, Omer Ali Bayraktar

**Author notes:** co-last & co-corresponding and.

## Abstract

The spatial organization of cell types in tissues fundamentally shapes cellular interactions and function, but the high-throughput spatial mapping of complex tissues remains a challenge. We present сell2location, a principled and versatile Bayesian model that integrates single-cell and spatial transcriptomics to map cell types *in situ* in a comprehensive manner. We show that сell2location outperforms existing tools in accuracy and comprehensiveness and we demonstrate its utility by mapping two complex tissues. In the mouse brain, we use a new paired single nucleus and spatial RNA-sequencing dataset to map dozens of cell types and identify tissue regions in an automated manner. We discover novel regional astrocyte subtypes including fine subpopulations in the thalamus and hypothalamus. In the human lymph node, we resolve spatially interlaced immune cell states and identify co-located groups of cells underlying tissue organisation. We spatially map a rare pre-germinal centre B-cell population and predict putative cellular interactions relevant to the interferon response. Collectively our results demonstrate how сell2location can serve as a versatile first-line analysis tool to map tissue architectures in a high-throughput manner.

## Introduction

The cellular architecture of tissues, where distinct cell types are organized in space, underlies cell-cell communication, organ function and pathology. Emerging spatial genomics technologies hold considerable promise for characterising tissue architecture, providing key opportunities to map resident cell types and cell signalling *in situ*, thereby helping guide *in vitro* tissue engineering efforts. Despite existing proof of concept applications^1–5^, it remains a challenge to define versatile and broadly applicable spatial genomics technologies and workflows. One reason is the enormous variation in tissue architecture across organs, ranging from the brain with hundreds of cell types found across discrete anatomical regions to immune organs with continuous cellular gradients and dynamically modified microenvironments. To create and map comprehensive tissue atlases, experimental and computational methods need to be aligned to cope with this variation and in particular, enable mapping numerous resident cell types across diverse and complex tissues *in situ*.

Strategies that generate coupled single-cell and spatially resolved transcriptomics offer a scalable approach to address these challenges. The key principle is to first identify resident cell types based on single-cell RNA-sequencing (scRNA-seq) from dissociated tissues, to then map the identified cell types to their tissue positions *in situ* based on spatial transcriptomic profiles^6^. Amongst spatial transcriptomic technologies, cyclic RNA imaging achieves single-cell resolution but is often limited to multiplexing a few hundred genes^7–9^. While longer cyclic imaging protocols can resolve more transcripts, imaging time limits the practical number of profiled genes, and hence the ability for unbiased cell type characterisation^10^. Alternatively, spatially resolved RNA-seq methods like Visium Spatial Transcriptomics^11^, HDST^12^ and Slide-sequencing^13^, where mRNAs are positionally captured from thin tissue sections using microarrays or bead-arrays, enable high-throughput data acquisition and rely on simple histology and molecular biology protocols.

In the above workflow, the step of mapping cell types using spatial transcriptomics poses several analytical challenges. First, spatial RNA-seq measurements (i.e. locations) combine multiple cell types as array-based mRNA capture currently do not match cellular boundaries in tissues. Thus, each spatial position corresponds to either several cell types (Visium, Tomo-Seq)^11,14^ or fractions of multiple cell types (Slide-Seq, HDST)^13^. Second, spatial RNA-seq measurements are confounded by different sources of variation as 1) cell numbers vary across tissue positions, 2) different cells and cell types differ in total mRNA content^15^, and 3) thin tissue sectioning captures variable fractions of each cell’s volume^16^. Computational approaches need to appropriately model and account for all of these factors.

Here, we present cell2location, a principled and versatile Bayesian model for comprehensive mapping of cell types in spatial transcriptomic data. Cell2location uses reference gene expression signatures of cell types derived from scRNA-seq to decompose multi-cell spatial transcriptomic data into cell type abundance maps. The model accurately maps complex tissues, including rare cell types and fine subtypes, and it identifies tissue regions and co-located cell types downstream in an automated manner. Besides, cell2location applies to different spatial RNA-seq technologies (Visium, Slide-Seq) and outperforms recently proposed methods in both accuracy and throughput. We demonstrate the power of cell2location to comprehensively map two model tissue types. In the mouse brain, we demonstrate a paired single nucleus and spatial transcriptomics workflow to discover novel regional astrocyte subtypes across the hypothalamus and thalamus. In the human lymph node, we resolve spatially interlaced cellular compartments, mapping rare B cell states and predicting putative cellular interactions relevant to the interferon response. These results present cell2location as a versatile tool for comprehensive mapping of tissue architecture.

## Results

### Cell2location: a Bayesian model for spatial mapping of cell types

Cell2location maps the spatial distribution of cell types by integrating single-cell RNA-seq (scRNA-seq) and multi-cell spatial transcriptomic data from a given tissue (Fig 1A). The first step of our model is to estimate reference cell type signatures from scRNA-seq profiles, for example as obtained using conventional clustering to identify cell types and subpopulations followed by estimation of average cluster gene expression profiles (Suppl. Methods, Fig S1). Cell2location implements this estimation step based on Negative Binomial regression, which allows to robustly combine data across technologies and batches (Suppl. Methods). In the second step, cell2location decomposes mRNA counts in spatial transcriptomic data using these reference signatures, thereby estimating the relative and absolute abundance of each cell type at each spatial location (Fig 1A, Fig S1).

**Figure 1.**
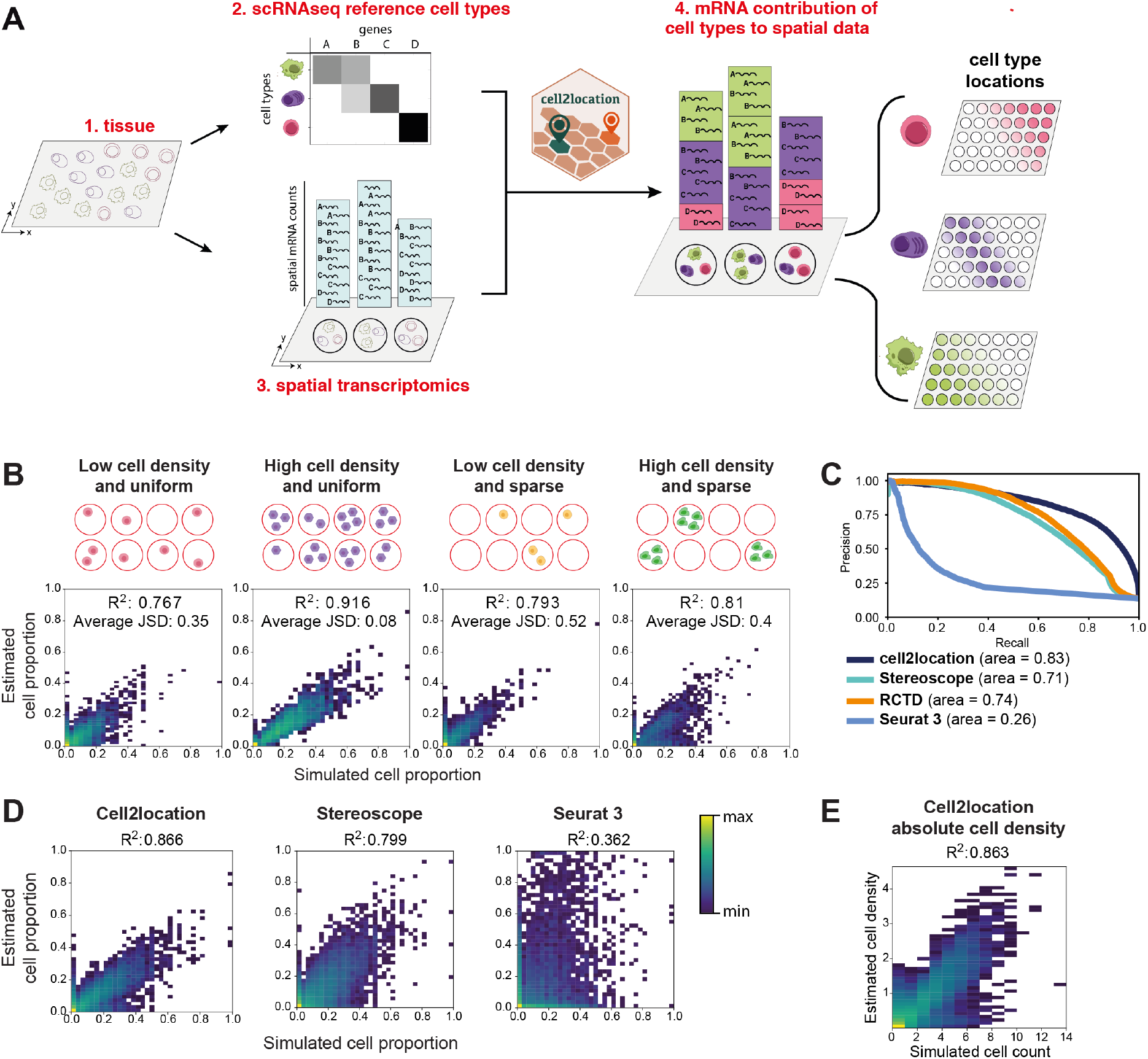
Cell2location model for spatial mapping of comprehensive cell type references. A. Overview of the spatial mapping approach and the workflow which are enabled by cell2location. From left to right: Single-cell RNA-seq and spatial transcriptomics profiles are generated from the same tissue (1). Cell2location takes reference cell type signatures derived from scRNA-seq and spatial transcriptomics data as input (2, 3). The model then decomposes spatially resolved multi-cell RNA counts matrices into the reference signatures, thereby establishing spatial maps of cell types (4). B. Model validation using simulated data. A benchmark dataset is constructed by combining cells drawn from 46 reference cell types (obtained from mouse brain scRNA-seq, Fig 2B) according to a synthetically generated cell type abundance map. Top: 4 alternative cell type abundance patterns considered for data simulation. Bottom: 2D histogram plots, displaying the concordance between simulated (X-axis) and estimated (Y-axis) cell-type proportions across 2,000 locations and cell types (Methods). Colour denotes 2D histogram count (50 bins along both X- and Y-axis). R2 denotes Pearson correlation and JSD denotes Jensen–Shannon divergence. C. Assessment of cell2location and alternative methods for detecting locations with non-zero cell abundance across all four cell type abundance patterns (2,000 locations, 46 cell types). Shown are precision-recall curves for considered methods with the corresponding areas under the curve stated in the legend. D. Assessment of cell2location and alternative methods for estimation of cell-type proportions across all four cell type abundance patterns (2,000 locations, 46 cell types). Shown are 2D histogram count plots (colour, 50 bins along both X- and Y-axis) between simulated (X-axis) and estimated (Y-axis) cell-type proportions for three alternative methods (left to right). E. Assessment of absolute cell abundance estimates obtained by cell2location. Shown is a 2D histogram count plot (colour, 50 bins along both X- and Y-axis) between simulated (X-axis) and estimated (Y-axis) cell abundance.

Cell2location is implemented as an interpretable hierarchical Bayesian model, thereby providing principled means to account for model uncertainty, (2) accounting for linear dependencies in cell type abundances, (3) modelling differences in measurement sensitivity across technologies, and (4) accounting for unexplained/residual variation by employing a flexible count-based error model. Finally, (5) cell2location is computationally efficient, owing to variational approximate inference and GPU acceleration. For full details and a comparison to existing approaches see Suppl. Methods. The cell2location software comes with a suite of downstream analysis tools, including the identification of groups of cell types with similar spatial locations.

To validate cell2location, we initially used simulated data that reflects diverse cell abundance and spatial patterns (Fig 1B). Briefly, we simulated a spatial transcriptomics dataset with 2,000 locations, based on reference cell-type annotations obtained from a mouse brain snRNA-seq reference dataset including 46 cell types (Methods). Multi-cell gene expression profiles at each location were derived by combining cells drawn from different reference cell types, using one of four cell abundance patterns with variable density and sparsity distribution that mimics the patterns observed in real data (Fig 1B; Methods). Assessing the concordance between estimated and simulated (true) cell proportions, we observed that cell2location mapped cell types with high accuracy across all considered cell abundance patterns (Fig 1B), including low abundant and sparsely located cell types.

Next, we compared cell2location to recently proposed alternative methods for the inference of relative cell-type abundance from spatial transcriptomics (Stereoscope^1^, Seurat^2^, RCTD^4^, NNLS (Autogenes)^17,18^, SPOTlight^3^, Fig S2). We found that cell2location was substantially more accurate in detecting the presence/absence of cell types across locations (Fig 1C, Fig S2), but the model also yielded more accurate estimates of relative cell type abundances (Fig 1D). Additionally, we note that cell2location is computationally more efficient than related model-based approaches for this task (Stereoscope: 14.8 hours on GPU versus cell2location: 8.66 minutes on GPU; Suppl. Methods). SPOTlight did not yield accurate results on the simulated data (Fig S2) and therefore we did not consider the model in our comparison. Finally, we note that unlike existing methods, cell2location not only provides estimates of relative cell type fractions but additionally estimates absolute cell type abundance, which can be interpreted as the number of cells that express a reference cell type signature at a given location, which again were highly concordant with the simulated ground truth (Fig 1E, Fig S3).

Collectively, these results support that cell2location can accurately estimate cell abundance across diverse cell types.

### Cell2location accurately maps mouse brain cell types

To illustrate how cell2location enables mapping a highly complex tissue, we examined the adult mouse brain, a tissue that contains highly diverse neuronal and glial cell types across stereotyped anatomical regions. We generated matched single nucleus (sn) and Visium spatial RNA-seq (10X Genomics) profiles of adjacent mouse brain sections that contain multiple regions from the telencephalon and diencephalon (Fig 2A). To assess the biological and intra-organ technical variation in spatial mapping, we assayed two mouse brains and serial tissue sections from each brain (total of 3 and 2 matched sections from two animals, respectively, and an extra section for snRNA-seq), creating a rich multi-modal and replicated transcriptomic dataset.

**Figure 2.**
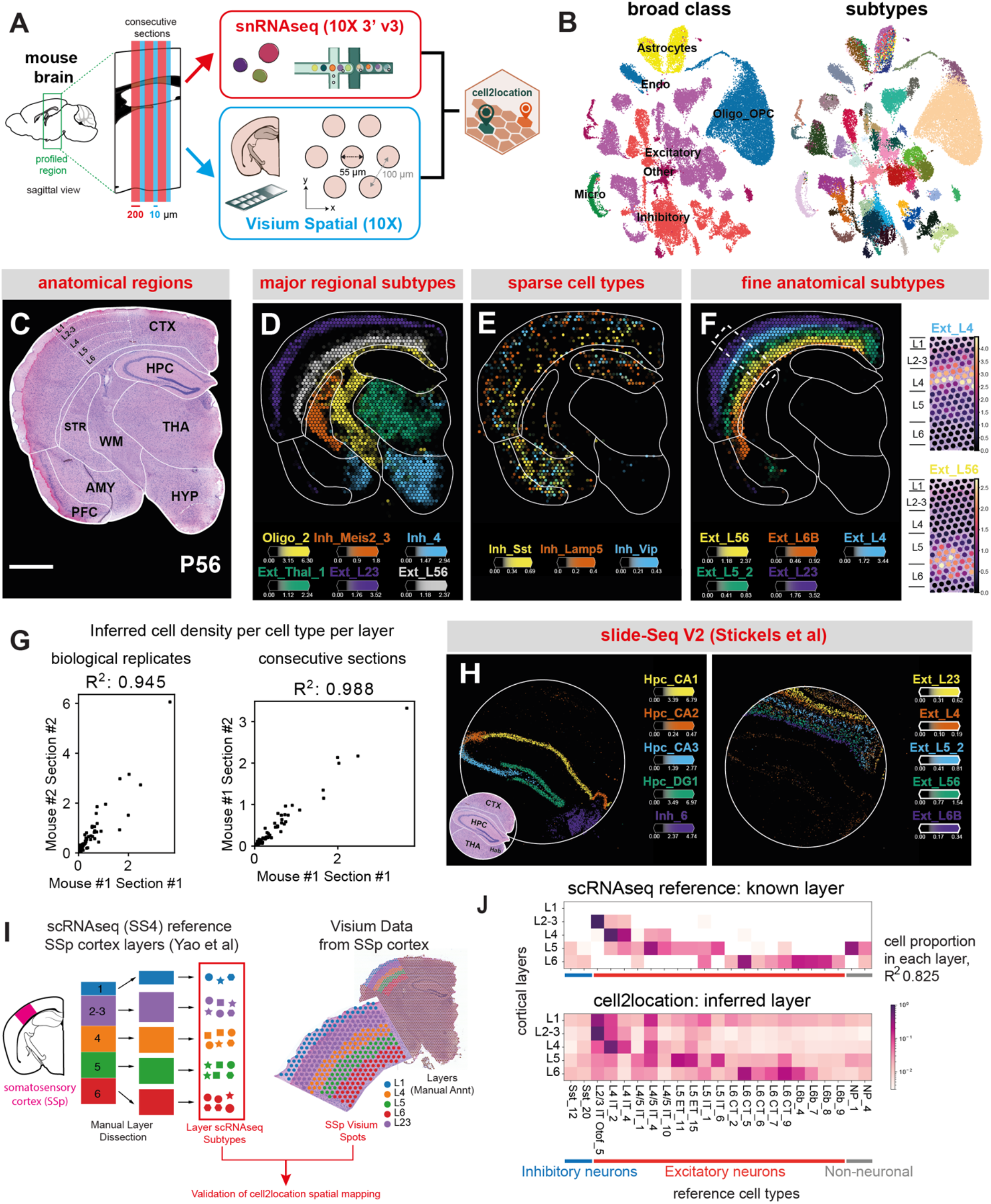
Comprehensive and accurate spatial mapping of cell types in the mouse brain. A. Experimental workflow. Adjacent tissue sections from two mice were used to generate single nucleus and spatial data, across a total of five pairings of Visium and snRNA-seq respectively (and an extra section for snRNA-seq). B. UMAP representation (X- and Y-axis) of broad cell classes (left) and 59 cell subtypes (right), as identified by Louvain clustering of the integrated snRNA-seq dataset across sections from (a). C. Mouse brain regions present in the assayed. H&E image of the tissue section from a postnatal day 56 (P56) brain and the outlines of brain regions are shown. Scale bar: 1 mm. D. Estimated cell densities (colour intensity) of major cell types (colour) across regions. E. Estimated cell densities (colour intensity) of sparse inhibitory neurons (colour). F. Estimated cell densities (colour intensity) of cortical excitatory neurons (colour). Small panels show the estimated cell density (colour) for two select subtypes on the region indicated with the white bounding box. G. Consistency of cell2location cell density estimates between replicate slides (X- and Y-axis), either considering replicate animals (left) or serial sections from the same animal (right). Considered are cell density estimates across 59 reference cell types across cortical layers; R2 denotes Pearson correlation. H. Spatial mapping in the Slide-Seq V2 data. Left: Estimated cell densities (colour intensity) of hippocampal neuron types (Hpc subtypes) and a thalamic habenular neuron type (Inh_6). Right: Estimated cell densities (colour intensity) of cortical excitatory neurons. I. Quantitative validation workflow based on scRNA-seq from manually dissected layers. Cortical layer neurons were mapped to SSp cortex region. Cell2location cell mapping accuracy was assessed by comparing estimated cell-type proportions for each subtype in each layer to estimates based on the scRNA-seq reference of neurons from manually dissected layers (Yao et al, 2020). J. Heatmap of relative cell subtype proportion (within each layer; colour, log10 scale) in each cortical layer in Allen reference (top) and the map of those cell types in Visium using cell2location (bottom).

We pooled snRNA-seq data across replicate animals and tissue sections (40,532 nuclei) and applied a conventional scRNA-seq analysis workflow followed by Louvain clustering to define reference cell-type signatures (Methods). This identified 59 cell clusters, which were well represented across replicate animals and sections (Fig S4). 43 clusters could be annotated as known putative cell types (Fig 2B, Fig S5, Methods), including excitatory or inhibitory neuron or glia subtypes, defined by markers from prior literature^15,19,20^. Additionally, we identified 6 previously known and 4 novel astrocyte subtypes (see next section). Importantly, our snRNA-seq dataset included cell types from all profiled brain areas (Fig S5), thus providing a regionally comprehensive and high-quality cell type reference for the analysis of the paired spatial data.

We applied cell2location to map these reference cell-type signatures to spatial locations of the mouse brain Visium data across replicate animals and sections (Fig S6). The resulting cell-type mapping was highly consistent with anatomical locations expected from prior literature (Fig 2C-F, Supp. Files 1-6). First, cell2location mapped abundant neuroglial cell types to their expected major brain regions, such as excitatory neurons of the cortex and thalamus^21^, inhibitory neurons of the striatum and hypothalamus, and oligodendrocytes of the white matter (Fig 2D)^20^. Second, cell2location mapped rare cortical inhibitory neurons to sparse locations, also as expected (Fig 2E)^22,23^. Third, cell2location spatially resolved fine anatomical subtypes within brain regions, such as excitatory neuronal subtypes across cortical layers and hippocampal divisions (Fig 2F, Fig S6E)^15,20^. To globally assess cell-type mapping across the full cell type reference, we used the output of cell2location across all 59 reference cell types and defined tissue regions based on clustering of locations with similar cell type abundance (Methods). This identified regions that correspond to anatomical brain structures, with the reference cell types mapping to the expected tissue regions (Fig S7). Finally, the cell2location estimates across manually annotated cortical layers were highly concordant across adjacent tissue sections and replicate animals (Fig 2G, Fig S8).

Next, to quantitatively assess the spatial cell-type mapping by cell2location, we used scRNA-seq (Smart-seq V4) datasets sampled from 5 manually dissected cortical layers and the hippocampus^24^ (Fig 2I). We generated corresponding reference signatures of 121 cell types, derived from the somatosensory cortex (n=98), as well as cell types that are exclusively present in the hippocampus (n=23). When using cell2location to map these cell types to the somatosensory cortex in our Visium data, the model almost exclusively mapped the cell types from the corresponding brain region (Fig S9), indicating high mapping specificity. Additionally, we selected the 23 most prevalent somatosensory cell types and assessed the consistency between relative cell-type abundance estimates across the five annotated layers estimated by cell2location and derived from scRNA-seq study^24^, finding a high overall concordance (Fig 2J, Fig S9C).

Finally, to demonstrate the versatility of cell2location across spatial transcriptomic technologies, we mapped our mouse brain snRNA-seq reference to a Slide-Seq dataset of the mouse brain that offers 10-micron spatial resolution^25^. Cell2location resolved cell types in Slide-Seq spatial data including fine anatomical subtypes across hippocampal divisions and cortical layers (Fig 2H, Fig S10A, Suppl. Files 7,8). Moreover, cell2location revealed that 50% of Slide-Seq “beads” (i.e. 10-micron diameter spatial locations) contained more than 1 cell type (Fig S10B), consistent with the previous observations^4^.

Collectively, our results demonstrate that cell2location maps diverse brain cell types with high accuracy and reproducibility; and that the method is broadly applicable to spatial transcriptomic technologies.

### Novel regional astrocyte subtypes

Next, we assessed cell2location for mapping closely related cell types in the mouse brain. We focused on the heterogeneity of astrocytes, the largest class of glial cells that intimately regulate neuronal circuit development and function^26^. While astrocytes were long considered a homogeneous cell type, recent evidence suggests that astrocytes are regionally diversified across the brain^26^, including fine anatomical divisions such as cortical layers^27–29^. However, the extent of the regional heterogeneity of astrocytes is largely unknown, as existing studies have primarily compared select brain areas^30,31^ or described broad astrocyte subtypes across major brain divisions (e.g. telencephalon versus diencephalon)^20,32^.

To identify regionally enriched astrocytes, we annotated and spatially mapped refined reference signatures of astrocyte subpopulations. Briefly, we considered 3,013 cells annotated as astrocytes from the initial analysis (Fig 2B), as identified based on the expression of canonical markers (e.g. *Aldoc*, *Slc1a3*). We then applied BBKNN^33^ to refine the integration of these cells across all 6 snRNA-seq datasets (Methods), followed by Leiden clustering, which identified 17 astrocyte subclusters (Fig S11). Next, we performed an initial mapping with cell2location and merged clusters that were not sufficiently distinct in their location and marker genes (Fig S12; Methods). This resulted in 10 molecularly and spatially distinct astrocyte subtypes (Fig 3A-D), which we considered for further analysis.

**Figure 3.**
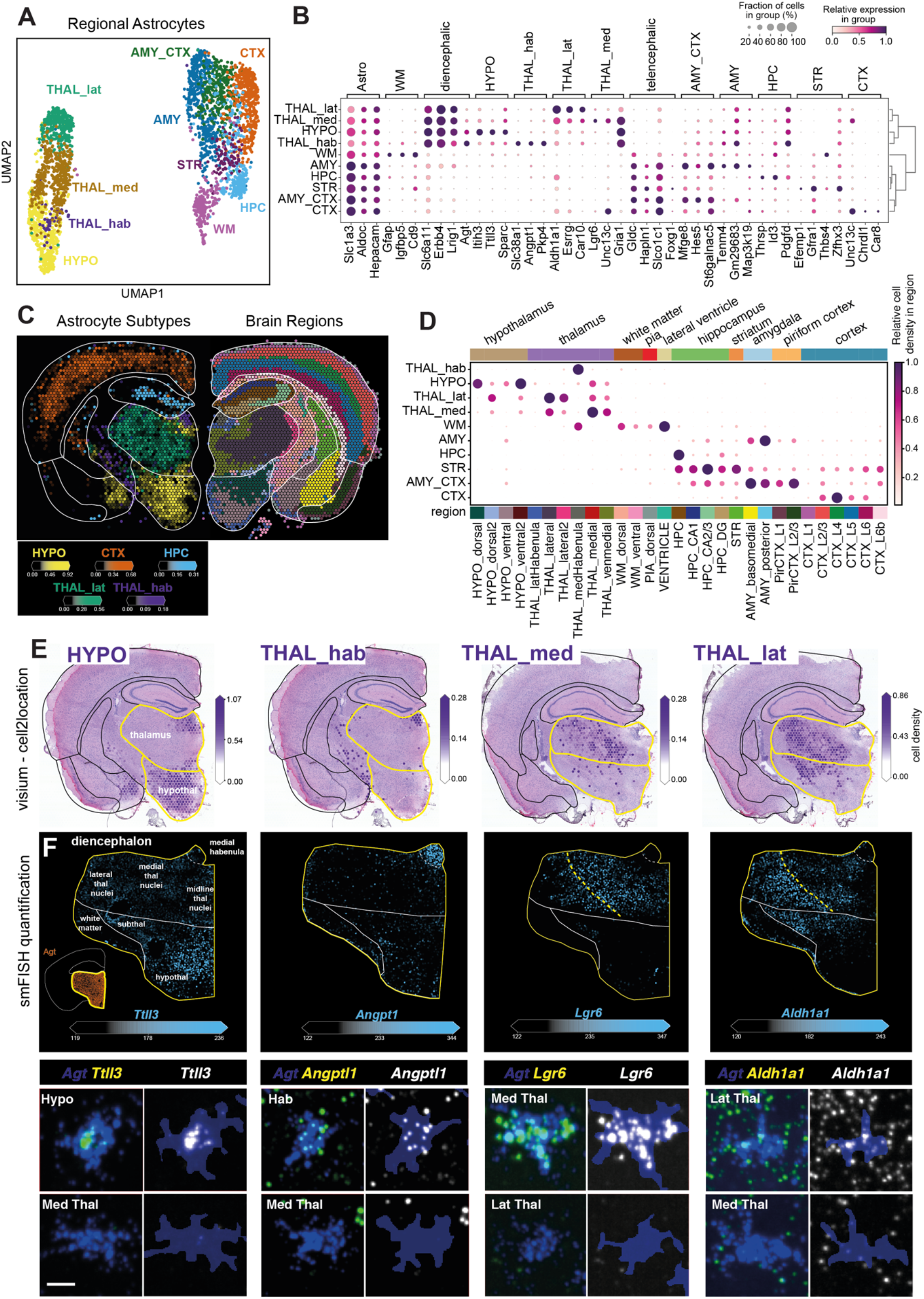
Spatial discovery of regional astrocyte subtypes. A. UMAP representation (X- and Y-axis) of 10 molecularly and spatially distinct astrocyte subtypes (colour) identified in our snRNA-seq data. B. Astrocyte subtype markers. Dot size corresponds to the fraction of cells in each astrocyte subtype cluster (rows) that express a given marker gene (count > 0; columns). Dot colour denotes the relative expression of each gene in each cluster, normalised by the maximum values for each gene (scaled between 0 and 1). Marker genes are grouped by subtypes. C. Estimated cell densities (colour intensity) of selected regional astrocyte subtypes (left hemisphere) and brain regions as identified using clustering of locations (right hemisphere). D. Estimated cell densities of astrocyte subtypes across brain regions (as in C). Dot plot, with dot size and colour corresponding to the relative cell density for each astrocyte subtype (rows) across regions (columns), scaled between 0 and 1 by normalising by the maximum values for each subtype. Colour bars denote broad brain areas (top) and subregion (bottom, matching panel C). E. Estimated cell densities (colour intensity) of 4 diencephalic astrocyte subtypes. Thal_med and Thal_lat are shown from a better representative section from the same mouse brain. F. smFISH validation. (Top panels) Small map shows locations of diencephalic astrocytes segmented based on Agt smFISH. Large maps show quantified subtype marker expression (colour intensity) at a single astrocyte level (dots). Yellow dashed line indicates the medial boundary of high Aldh1a1 expression. (Bottom panels) Close-up images of diencephalic astrocyte subtypes. First columns show smFISH for astrocyte marker Agt and subtype markers. Second columns show astrocyte segmentation masks (blue) transparently overlaid on subtype genes. Scale bar: 10 microns.

We identified grey matter astrocyte subtypes that were spatially enriched in the thalamus (THAL), hypothalamus (HYPO), cortex (CTX), amygdala (AMY), hippocampus (HPC) and striatum (STR) as well as white matter (WM) astrocytes (Fig 3C, D). These subtypes expressed expected regional astrocyte markers^20^, such as *Foxg1* in the telencephalon (CTX, AMY, HPC, STR) and *Slc6a11* and *Agt* in the diencephalon (THAL, HYPO) (Fig 3B). Moreover, they stratified to finer regional subtypes based on the expression of new subtype-specific genes such as *Angpt1* and combinatorially expressed markers such as *Gria1* (Fig 3B, see below).

We focused on diencephalic astrocyte subtypes as only one type of grey matter astrocyte subtype has been described in this brain area to date^20^. Notably, our data revealed 4 regionally distinct grey matter astrocyte subpopulations across the diencephalon (Fig 3E, Fig S13). HYPO astrocytes were mainly located in the hypothalamus and, to a lesser extent, in the midline thalamic nuclei and the lateral habenula, and were marked by *Ttll3* expression. Three THAL subtypes occupied different regions of the thalamus. THAL_hab astrocytes, a highly rare cell population in our reference (Fig S4), were restricted to the medial habenula and ventral white matter tracts, and specifically expressed *Angpt1* (Fig 3E). THAL_med astrocytes were enriched in the medial thalamic nuclei, and were marked by *Lgr6* expression. Finally, THAL_lat astrocytes, while present throughout the thalamus and the subthalamus, were most prevalent in ventrolateral thalamic nuclei, and were marked by elevated *Aldh1a1* expression compared to other subtypes (Fig 3E).

We validated the identity and distribution of diencephalic astrocyte subtypes using single molecule fluorescent *in situ* hybridization (smFISH)^27^. We used *Agt* smFISH to specifically label and segment astrocytes throughout the diencephalon^20^ (Fig S14, Fig S15), then quantified the expression of co-stained subtype markers at single astrocyte resolution (Fig 3F). *Ttll3* showed the highest expression in hypothalamic astrocytes, while *Angpt1* was highly enriched in astrocytes of the medial habenula. *Lgr6* expression peaked in astrocytes in medial thalamic nuclei while *Ald1ha1* expression was enriched in the lateral thalamus in a complementary manner, both genes did not express in habenular astrocytes. These results validate diencephalic astrocyte subtypes and demonstrate that cell2location can spatially map closely related cell types.

### Mapping the cellular compartments of the human lymph node

To further explore the application of cell2location to complex tissue architectures, we applied the model to spatially map the human lymph node. Unlike the mouse brain, the human lymph node is characterised by dynamic micro-environments with many spatially interlaced cell populations. We analysed a publicly available Visium dataset of the human lymph node from 10X Genomics^34^, spatially mapping a comprehensive atlas of 34 reference cell types derived by integration of scRNA-seq datasets from human secondary lymphoid organs composed of 73,260 cells^35–37^ (Fig 4A-B; Methods).

**Figure 4.**
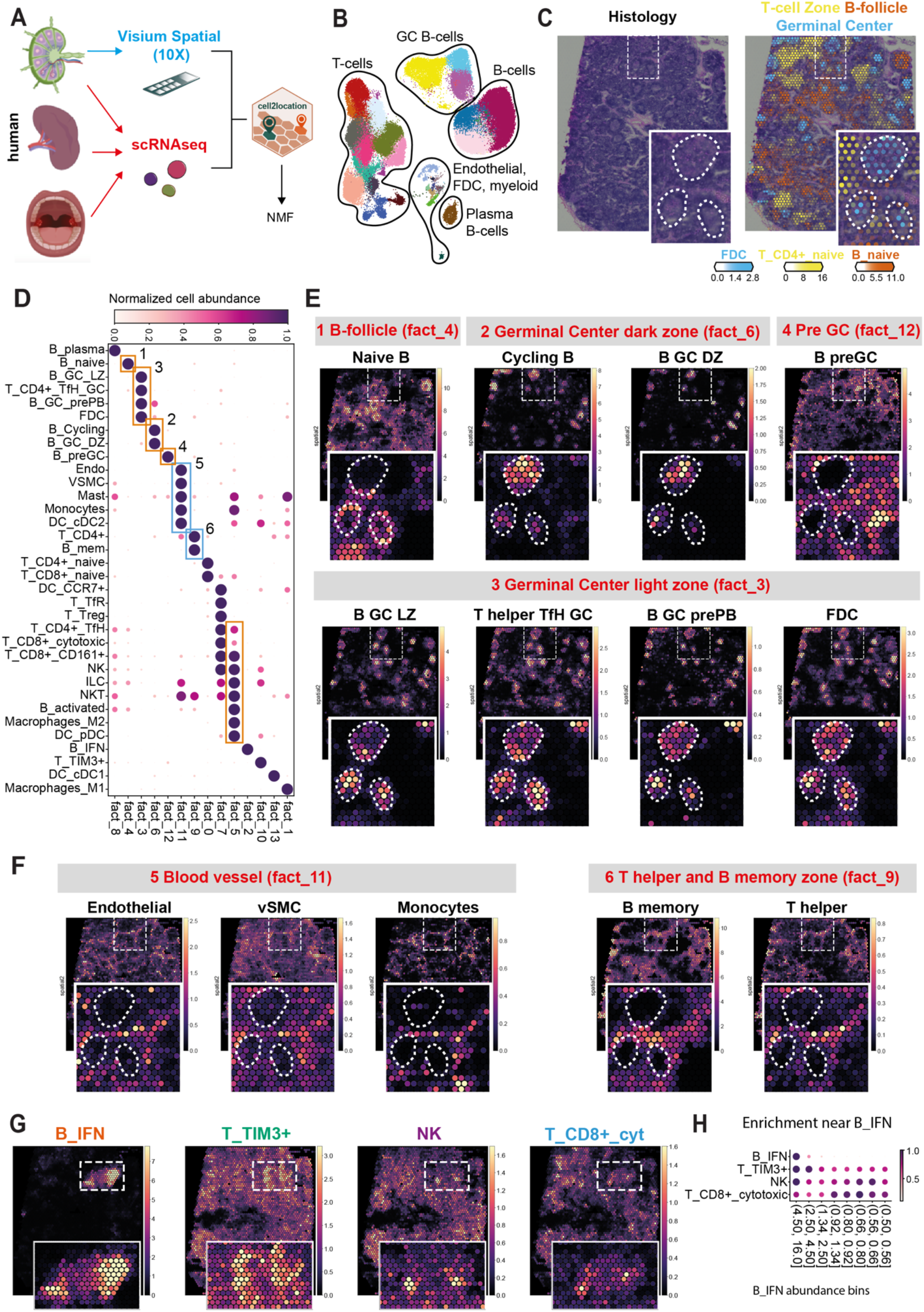
Mapping the cellular compartments of the human lymph node. A. Cell2location analysis workflow. B. UMAP representation (X- and Y-axis) of 34 immune and non-immune cell subtypes (colour), combining scRNA-seq data from 3 studies used to define reference gene expression signatures of cell types. C. Major tissue zones of the lymph node, including T-cell and B-cell zones, and GC reactions with FDCs. Left: H&E image of the tissue section. Right: Estimated cell density (colour intensity) of cell types (colour) shown over the H&E image of the lymph node sample. D. Identification of lymph node cell compartments using non-negative matrix factorization. Shown is a dot plot of the NMF weights of cell types (rows) across NMF components (columns), which correspond to cell compartments (normalized across components per cell type by dividing by maximum values). E. Location patterns of cell types that are associated with highlighted cell compartments from Fig 4D. Spatial plots (X, Y-axis) show cell density (colour) for each cell type (subpanels). Dashed boxes highlight the tissue regions shown in inset panels. Dashed circles indicate GCs. F. Location patterns of the blood vessel and perivascular cell types that are associated with highlighted cell compartments from Fig 4D. Plotted as in above. G. Co-localisation of cytotoxic NK, T cells and exhausted T cells with B cells expressing interferon response genes (B IFN). Spatial plots show cell density (colour) for each cell type (subpanels). Dashed lines indicate regions shown in inset boxes. H. Quantifying enrichment cytotoxic NK, T cells and exhausted T cells in proximity of B IFN cells. Dot plots show average cell density (colour and size) of each cell type (rows) across B_IFN abundance quantile bins (columns).

Histological examination of the lymph node Visium sample revealed multiple germinal centres (GCs). Correspondingly, following mapping with cell2location, the expected major regions within the lymph node tissue could be readily identified, including T-cell and B-cell zones, and GC reactions with follicular dendritic cells (FDC) (Fig 4C, right panel, Suppl. Files 9-10).

Next, we set out to explore the downstream analysis of cell2location mapping results to identify the spatial co-occurrence of cell types, which can aid the discovery of tissue organisation and prediction of cellular interactions. We performed non-negative matrix factorization (NMF) of the cell type abundance estimates from cell2location (Fig 4D). Similar to the known benefits of NMF applied to conventional scRNA-seq^38,39^, this analysis provides an additive decomposition of the spatial cell type densities into components that correspond to groups of co-localised cell types across locations. This matrix decomposition naturally accounts for the fact that multiple cell types and microenvironments can co-exist at the same Visium locations (Fig S16), while sharing information across tissue areas (e.g. individual germinal centres).

We observed that the groups of cell types identified by NMF corresponded to known functionally relevant cellular compartments of the lymph node (Fig 4D). Furthermore, the NMF output enabled the dissection of the spatial compartments of B cell maturation. Specifically, naive B cells were mapped to a B-follicle zone distinct from germinal centres (Fig 4E, panel 1). Before GC entry or formation, B cells are activated through acquiring antigen from antigen-presenting cells such as macrophages and dendritic cells - these interacting cell types were consistently co-located (Fig 4D, factor #5). We also identified the GC dark zone where B cells clonally expand and proliferate (Fig 4E, panel 2) as well as the GC light zone where they undergo selection by T follicular helper cells and FDCs to differentiate into antibody-producing plasmablasts (Fig 4E, panel 3).

Beyond confirming known biology, the NMF analysis enabled the fine-grained spatial mapping of a rare and putatively novel secondary activation B cell state, which has been recently reported as primed to undergo class switch recombination preceding the GC entry^37,40^. Our data indicate that this population (termed here “preGC”) occupies a different tissue compartment from the upstream activated cells and downstream GC cells (Fig 4E, panel 4), suggesting it is more likely to initiate new GC reactions rather than joining existing ones. Finally, our analysis was capable of distinguishing blood vessel zones containing endothelial and vascular smooth muscle cells (Fig 4F, panel 5) from the adjacent perivascular zone enriched in T helper and B memory cells (Fig 4F, panel 6).

We noted that rare B cells with a distinct interferon response gene signature (B IFN) were spatially segregated from all other B-cell and germinal centre zones (Fig 4G), suggesting they might represent virally infected B-cells or autoreactive B cells^41^. Spatial enrichment indicated the recruitment of cytotoxic T cells and NK cells to this zone (Fig 4H), including TIM3+ T cells expressing markers of T cell exhaustion and acute viral infection^42,43^. Hence, we hypothesise that these cytotoxic cells are responding to a viral infection in B-cells and are becoming exhausted after attempting to contain the infection.

Taken together, our results show that cell2location can map complex tissues with spatially interlaced cell types and aid biological interpretation by identifying putatively interacting cell types.

## Discussion

Cell atlassing efforts are generating increasingly complex single cell and spatial transcriptomic datasets of diverse tissues. Here, we presented cell2location, a Bayesian model that integrates single-cell and spatial transcriptomics to map cell atlases in a scalable manner. The model can cope with complex tissues, handling large numbers of cell types including subtypes characterized by subtle transcriptional differences. Owing to a number of technical innovations, we find that cell2location is more accurate than existing methods, enabling efficient spatial mapping of tens to hundreds of reference cell types.

We demonstrated that cell2location can integrate scRNA-seq and coarse spatial resolution transcriptomic data such as Visium to map tissue atlases with high resolution. Applied in this workflow, cell2location can accurately pinpoint rare reference cell types like habenula-specific astrocytes to few locations in spatial data. We envision that our approach will be applied to spatially map the “first generation” cell type atlases of large tissues, such as the whole mouse brain^20,44^, at moderate cost and high throughput.

We show that cell2location spatial mapping can aid the discovery of new cell types, as illustrated by 4 regionally distinct astrocyte subtypes we describe in the mouse diencephalon. We can only speculate about the regionalized functions of these astrocytes. Astrocyte-neuron signalling in the lateral habenula regulates neuronal circuits implicated in depression^45^. Similarly, diencephalic astrocytes could shape the activity of local neuronal circuits, such as the lateral thalamic circuitry for sensory relay or medial thalamic circuits involved in attention control^46^.

In an application to the human lymph node, we demonstrated that cell2location can disentangle spatially intermixed cell types. We have combined cell2location with NMF to provide a principled strategy for resolving tissue compartments with stereotyped cell composition, such as germinal centre zones. This approach can elucidate the wiring diagram of cell types in complex tissues by spatially mapping putatively interacting cell types and help prioritise interactions predicted from single cell data^47^.

We expect future developments of cell2location and related models to spatially map large tissues, scaling up to hundreds of thousands or millions of spatial locations. Future extensions will also provide adaptations to specific spatial technologies, thereby accounting for differences in the noise characteristics such as background probe binding or variable capture areas (e.g. Nanostring^48^, LCM-RNA-seq^49^).

A major future frontier for cell atlas projects is the 3D mapping of tissues at cellular resolution. The spatial mapping approach described here can lay the foundations for 3D tissue atlases. Single cell, spatial RNAseq and imaging can be applied to consecutive tissue sections across an organ, integrating data from all technologies to build a multi-scale atlas. Regional cell type maps generated with cell2location can also guide gene selection for intact tissue volume imaging^50^, leading to 3D models of cells in their native tissue context with whole transcriptome information.

In summary, we have here introduced a versatile and flexible approach for spatial mapping of tissues. We expect that cell2location will have diverse applications in mapping tissue architecture across development, health and disease. Spatial cell type maps coupled with cell interaction analysis can help decipher developmental processes such as cellular differentiation and how it is regulated by extrinsic cues. These developmental cues can be recapitulated *in vitro* to refine tissue engineering approaches. Similarly, understanding how tissue architecture is perturbed in response to disease can enrich our understanding of pathological processes and aid the design of therapeutic interventions.

## Methods

### Cell2location model

For a complete derivation of the cell2location model, please see supplementary computational methods. Briefly, cell2location is a Bayesian model, which estimates absolute cell density of cell types by decomposing mRNA counts *d_s,g_* of each gene *g* = {1, . ., *G*} at locations *g* = {1, . ., *G*} into a set of predefined reference signatures of cell types *g_fg_*.

#### Estimation of cell type reference signatures from scRNA-seq

Given cell type annotation for each cell, the corresponding reference cell type signatures *g_fg_*, which represent the average mRNA count of each gene *g* in each cell type *f* = {1, . ., *F*}, can be estimated using a negative binomial regression model, which allows for combining data across batches and technologies (Suppl. methods).

#### Cell2location model

An untransformed spatial expression count matrix *d_s,g_* is used for input, as obtained from the 10X SpaceRanger software (10X Visium data). Cell2location models the elements of *d_s,g_* as Negative Binomial (NB) distributed, given an unobserved gene expression level (rate) *μ_s,g_* and a gene-specific over-dispersion *α_g_*:

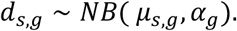

The expression level of genes *μ_s,g_* in the mRNA count space is modelled as a linear function of expression signatures of reference cell types:

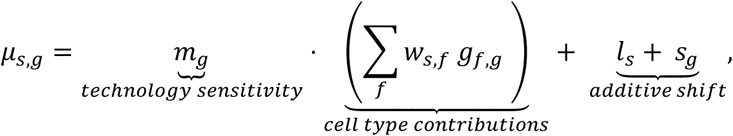

where, *w_s,f_* denotes regression weight of each reference signature *f* at location *s*, which can be interpreted as the number of cells at location *s* that express reference signature *f*;*m_g_* is a gene-specific scaling parameter, which adjusts for global differences in sensitivity between technologies; *l_s_* and *s*_g_ are additive variables that account for gene- and location-specific shift, such as due to contaminating or free-floating RNA.

To account for the similarity of location patterns across cell types, *w_s,f_* is modelled using another layer of decomposition (factorization) using *r* = {1, . ., *R*} groups of cell types, that can be interpreted as cellular compartments or tissue zones (Suppl. Methods). Unless stated otherwise, *R* is set to 50.

While the scaling parameter *m_g_* facilitates the integration across technologies, it leads to non-identifiability between *m_g_* and *w_s,f_*, unless the informative priors on both variables are used. To address this, we employ informative prior distributions on *w_s,f_* and *m_g_*, which are controlled by 4 used-provided hyper-parameters: 1) 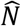, the expected number of cells per location; 2) 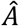, the expected number of cell types per location; 3) 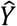, the expected number of co-located cell type groups per location; 4) mean μ and variance *σ*^2^ that define hyperprior on gene-specific scaling parameter *m_g_*, allowing the user to define prior beliefs on the sensitivity of spatial technology compared to the scRNA-seq reference. For guidance on selecting these hyper-parameters see Suppl. Methods (Section 1.3) and the methods section on spatial mapping of the mouse brain cell types below.

Approximate Variational Inference is used to estimate the parameters, implemented in the pymc3 framework ^51^, which supports GPU acceleration. For full details see Suppl. Methods.

#### Note on constructing reference cell type data

It is important to aim for a comprehensive and detailed cell type reference which includes the cell types and sub-populations that are present *in-situ*, for example, by generating a paired snRNA-seq reference from the same tissue sample. However, imperfect matching of cell populations is often acceptable (see Fig 4 as an example). In such instances, the stability of the model fit, which can be assessed multiple random restarts, can be used as diagnostics (see Suppl. Methods).

### Mouse brain tissue processing for matched snRNA-seq and Visium

Mouse work was carried out at the Wellcome Sanger Institute in accordance with UK Home Office regulations and UK Animals (Scientific Procedures) Act of 1986 under a UK Home Office license, which were regularly reviewed by the institutional Animal Welfare and Ethical Review Body.

Brains of wild-type adult C57BL/6 mice (postnatal day 56, 1 female and 1 male) were dissected, snap frozen, embedded in optimal cutting temperature compound (Tissue-Tek) and stored at −80°C. Brain hemispheres were cryosectioned at −20°C using a cryostat (Leica, CM3050S). To assess tissue quality, RNA was extracted from test tissue sections from each animal using the RNeasy Pico Kit (Qiagen) and yielded high RIN values (9.6 and 9.7) on an Agilent Bioanalyser, indicating high RNA quality.

For matched single nuclei and Visium RNA-seq experiments, brain hemispheres were cryosectioned to adjacent thick (200 μm) and thin (10 μm) coronal sections, respectively, and processed the same day. In total, four consecutive sets of thick and thin tissue sections were collected from each brain. Five sets of tissue sections yielded both good quality single nuclei and Visium data (three adjacent sections from mouse 1 and two sections from mouse 2) while one additional section from mouse 2 yielded good single nuclei; these were considered for analysis in this study.

### Single nucleus RNA-sequencing

Thick (200 μm) mouse brain sections were cryosectioned, dissected from OCT and kept in a tube on dry ice until subsequent processing. Nuclei were extracted from each section as described previously^52^. Briefly, nuclei were released from sections via Dounce homogenisation, Hoechst-stained, and isolated via fluorescence-activated cell sorting (FACS). Nuclei were then loaded into the 10X Chromium Single Cell 3′ Kit (v3) to obtain 3000-7000 nuclei per well, and library preparation was done per manufacturer’s protocol. Libraries were sequenced on an Illumina NovaSeq S4 system.

### Visium spatial transcriptomics

Thin (10 μm) mouse brain sections were cryosectioned and mounted directly onto separate capture areas on 10X Visium Spatial Gene Expression slides (beta product version). Processing was done per manufacturer’s protocols. Briefly, sections were methanol-fixed, hematoxylin and eosin (H&E)-stained, and imaged on a NanoZoomer 2.0 slide scanner (Hamamatsu). Sections were then permeabilized and further processed to obtain cDNA libraries that were quality controlled using the Agilent Bioanalyser. The cDNA libraries were sequenced on the Illumina HiSeq 4000 system, aiming 300 million raw reads per section with read lengths 28cy R1, 8cy i7 index, 0cy i5 index, 91cy read 2.

### Single nucleus data processing and cell-type annotation

Sequencing data were processed using 10X CellRanger version 3.0.2, aligned to mouse pre-mRNA genome reference version mm10 and mRNA count matrices were generated by adding intronic and exonic unique molecular identifier (UMI) counts for each gene in each cell. snRNA-seq counts were processed using standard Seurat V3 workflow without correcting batch effects between 6 individual samples. Specifically, 10X CellRanger output, expression matrices filtered to include droplets containing cells, was imported into the Seurat object. Cells were filtered using the following quality control criteria: number of detected genes per cell < 5000, fraction of nuclear-encoded mitochondrial genes < 0.10, doublet scores (scrublet^53^) < 0.2. The data was then subjected to normalisation with scale-factor of 10000 and log1p-transformed (Seurat::NormalizeData), scaled gene-wise by subtracting the mean and dividing by standard deviation (Seurat::ScaleData). Following that, 50 principal components were found (Seurat::RunPCA) and used as input to UMAP algorithm (Seurat::RunUMAP), which was also used for K-nearest-neighbour graph (KNN) construction with k=100 (Seurat::FindNeighbors). To generate a comprehensive cell annotation, joint Louvain clustering with resolution 7 was performed (Seurat::FindClusters) on all samples. Clusters were subsequently annotated using marker genes from the literature and existing mouse brain single cell reference datasets^15,19,20^. To annotate the regional identity of cell types, we used 1) known regional marker genes^15,19,20^, 2) the spatial expression patterns of these genes on *in-situ* hybridization data from the Allen Brain Atlas (https://mouse.brain-map.org/), 3) the regional localization of cell types estimated by cell2location (Fig 2, Suppl. Files 1-5). The marker genes of regional cell types reported in Fig S5 were obtained from Zeisel et al^20^ (http://mousebrain.org/downloads.html, L5_All.agg.loom file). The resulting clusters and cell annotations are reported in Fig 2B, Fig S4, Fig S5 and were used as input to cell2location.

### Visium data processing

10X Visium spatial sequencing data was aligned to mouse pre-mRNA genome reference version mm10 using 10X SpaceRanger and mRNA count matrices were generated by adding intronic and exonic reads for each gene in each location. The paired histology H&E images were processed using 10X SpaceRanger to select locations covered by tissue by aligning pre-recorded spot locations with fiducial border spots in the histology image. This allowed evaluating the correspondence between cell maps produced using our method and the known brain anatomy. This also enabled the quantification the number of nuclei in each spot using image segmentation as described in Suppl. methods and reported in Fig S6A-D. The histology image was used to manually annotate cortical layers in the primary somatosensory cortex (SSp) region using the lasso tool in the 10X Loupe browser.

### Constructing synthetic spatial transcriptomics data set

Simulated spatial transcriptomics data were generated by combining expression profiles of cells, drawn from each of the 46 reference cell types in the mouse brain snRNA-seq reference data, according to simulated abundance at 2000 locations. The snRNA data for the two most homogenous mouse brain snRNA-seq samples was split into the dataset used to generate the synthetic data (50% of cells) and the dataset used to evaluate cell2location and alternative approaches (50% of cells), similarly to the strategy proposed by Andersson et al^1^. The number of cell types present in each simulated location as well as their absolute abundance (the number of cells per location) were simulated according to four cell type sparsity and abundance patterns profiles - uniform and sparse, low and high average abundance (Fig 1B, see below). The specific parameters of each pattern were chosen to mimic those observed in real data (low cell count of each cell type, low total number of cells, high sparsity). The code used to generate simulated data can be obtained from https://github.com/emdann/ST_simulation/.

1. **Generating per cell-type sparsity and density patterns** (assemble_design tool).

a. **Uniform and low density:** 40% of cell types present in *N* ~ *Gamma*(*μ* = 2000. 0.8, *σ*^2^ = *μ* / 0.3) locations at density *D* ~ *Gamma*(*μ* = 0.8, *σ*^2^ = *μ*).
b. **Uniform and high density:** 10% of cell types present in *N* ~ *Gamma*(*μ* = 2000. 0.8, *σ*^2^ = *μ* / 0.3) locations at density *D* ~ *Gamma*(*μ* = 2.5, *σ*^2^ = *μ*).
c. **Sparse and low density:** 40% of cell types present in *N* ~ *Gamma*(*μ* = 2000. 0.1, *σ*^2^ = *μ* / 0.3) locations at density*D* ~ *Gamma*(*μ* = 0.8, *σ*^2^ = *μ*).
d. **Sparse and high density:** 10% of cell types present in *N* ~ *Gamma*(*μ* = 2000. 0.1, *σ*^2^ = *μ* / 0.3) locations at density *D* ~ *Gamma*(*μ* = 2.5, *σ*^2^ = *μ*). By following this procedure, sparsity and density parameters for each cell type were generated to produce an average number of cells per location close to 10, mimicking cell count observed by nuclear segmentation of the mouse brain histology images (Fig S6; Suppl. Methods).
2. **Assembling abundance of each cell type in each location** (assemble_composition tool). For each cell type, N random locations (N determined in step 1) are chosen from all 2000 locations, and for each of those positive locations, the number of cells M was generated by sampling from a Poisson distribution with a rate equal to the density parameter D selected in step 1. The cell number for remaining locations was set to 0.
3. **Assembling mRNA counts of each gene in each location.** Multi-cell mRNA count profiles for every gene in a given location were derived by combining M cells drawn from reference cell types in the snRNA-seq data (using M determined for every cell type and location in step 2).
4. **Applying gene-specific scaling to mimic the difference in sensitivity between technologies.** A dataset with lower sensitivity was generated by taking samples from the Poisson distribution with a rate equal to the mRNA counts in the synthetic dataset generated in step 4 multiplied by the gene-specific scaling parameter. This parameter L was randomly generated from *L*; ~ *Gamma*(α = 0.41885602, β = 0.9527432), with parameters selected to mimic those estimated by cell2location in the mouse brain data. Gene-specific L parameters were divided by 2 to mimic reduction in sensitivity.

### Validation of cell2location using synthetic data and comparison to alternative methods

Synthetic gene expression data were constructed as described above and were used to assess the performance of cell2location in comparison to alternative methods. Synthetic data came with ground truth numbers of cells and mRNA count of each cell type for each location, which were used to benchmark estimates obtained from alternative methods. The models were provided with the snRNA-seq reference data (evaluation subset, 8137 cells by 12281 genes, see above) as well as the synthetic mRNA count matrices for all locations.

#### Cell2location

Signatures of reference cell types were derived using a negative binomial regression model (Suppl. Methods) with default parameters, except the L2 penalty for overdispersion parameter: ‘gene_overdisp_weight’ = 0.1. Cell2location was used in the single-experiment mode with the following priors and parameters:

- Training iterations: 30000,
- Cells per location 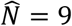; Cell types per location 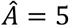; Co-located cell type groups per location 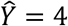;
- Prior on the difference in sensitivity between technologies: **μ** = 1/2, *σ*^2^ = 1/8;

A simplified version of the cell2location model, which does not account for linear dependencies in the abundance of cell types was considered for comparison, using otherwise identical hyperparameters (Fig S2).

#### Stereoscope^1^

This method utilises a similar model-based approach to estimating both the cell type signatures and the relative cell abundance across locations (see the comparison in Suppl. Methods). In the Stereoscope model (version 0.2), Negative Binomial distribution is parameterised using the gene- and cell type-specific log-odds parameter and gene-specific total count parameter. This custom signature estimation method is hard-coded into the workflow and both estimated parameters are used as input to the model for estimating relative cell abundance. Stereoscope was used with the following hyperparameters as suggested in the tutorial on GitHub (https://github.com/almaan/stereoscope/tree/f42a0a26305963e587b372c44094db108b26b6e8#3-analysis):

- Highly variable gene selection: 5000 genes
- Estimating expression signatures of cell types: 75000 iterations, 1000 mini-batch training size
- Estimating relative cell abundance: 75000 iterations, 1000 mini-batch training size

#### RCTD^4^

This method utilises a similar model-based approach to the estimation of relative cell abundance across locations (see the comparison in Suppl Methods). It estimates reference cell type signatures by computing expression average per cluster (hard-coded). RCTD was used in default mode (not doublet mode) with the hyperparameters selected as suggested in the tutorial on GitHub, including filtering down to 2763 genes based on log-fold-change (https://github.com/dmcable/RCTD/tree/7a895ed1666269784cc472fa0a6e4abbd3d2ccb2).

#### Seurat V3 (PCA, anchor method, MNN classifier)^2^

Seurat V3 workflow was used to project synthetic locations into the PCA space constructed using snRNA-seq reference data (Seurat V3 with anchor selection, Seurat::FindTransferAnchors), with subsequent transfer of cell type labels onto synthetic locations using a mutual nearest neighbour (MNN)-classifier (Seurat::TransferData). For the purpose of this comparison, the label transfer scores were interpreted as a relative cell abundance of each cell type in each location. Prior to integration both snRNA-seq reference and synthetic spatial datasets were subjected to SCTransform normalisation (Seurat::SCTransform)^54^. PCA method (with default 30 components) was used for integration (Seurat::FindTransferAnchors) as advised in Seurat package documentation and spatial mapping tutorial. All functions were used with default parameters.

#### SPOTlight^3^

SPOTlight uses a seeded topic model to find expression signatures of reference cell types and a modified non-negative least squares method (denoted NMFreg) to estimate relative cell abundance. The topic model is initialized in a way that guides the topics to resemble expression signatures of cell types. SPOTlight workflow was followed with recommended parameters (https://github.com/MarcElosua/SPOTlight/tree/fb5e7c7de3e6d0ac7618c4b1a55aea06e5f472ea).

Briefly, to improve computational efficiency the snRNA-seq data size was reduced by randomly selecting at most 100 cells per cell type (when a cell type is represented by less than 100 cells all cells are kept), and highly variable genes (5000 genes) were selected using Seurat::FindAllMarkers tool. Untransformed expression counts for both snRNA-seq reference and synthetic locations were used as input to ‘spotlight_deconvolution’ function in SPOTlight package with the following hyper-parameters: transf = “uv”, method = “nsNMF”, min_cont = 0.09.

#### Non-negative least squares (NNLS, autogenes)^17,18^

NNLS was used to estimate the absolute cell abundance of each cell type in each synthetic location given the reference signatures of cell types, which were derived by computing average expression of each gene across cells of each type. Untransformed expression counts of the synthetic locations were used as input to NNLS implementation in autogenes package. Similarly to cell2location, the output was normalized to produce relative cell abundance.

The estimates of relative cell abundance from all methods were compared to the ground truth using 2D histogram plots, Pearson R^2^ (numpy.corrcoef) and Jensen–Shannon divergence (scipy.spatial.distance.jensenshannon) and reported in Fig 1B, Fig 1D, Fig 1E, Fig S2C, Fig S3. In addition, the accuracy was assessed using precision-recall quantified on all 46 cell types jointly (micro-average). Relative cell abundance was used as a predictor and cell count > 0 as a gold standard label. Precision-recall results are reported in Fig 1C, Fig S2A.

### Spatial mapping of mouse brain cell types to 10X Visium data

Expression signatures of 59 mouse brain reference cell types (Fig 2B, see above) were derived using a negative binomial regression model (Suppl. Methods). Mouse brain 10X Visium data was processed as described above to untransformed mRNA counts (filtered to 12,809 genes shared with scRNA-seq, 14,968 locations), which were used as input to cell2location using the following parameters:

- Training iterations: 30000.
- Cells per location 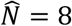, estimated based on comparison to histology image; Cell types per location 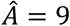, assuming that most cells in a given location belong to a different type and that many locations contain cell processes rather than complete cells; Co-located cell type groups per location 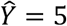, assuming that very few cell types have linearly dependent abundance patterns, except for the regional astrocytes and corresponding neurons such that on average about 2 cell types per group are expected 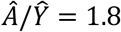.
- Prior on the difference in sensitivity between technologies: *μ* = 1/2, *σ*^2^ = 1/8, reflecting that the average total mRNA count in snRNA-seq is about 2 times greater than the average total mRNA count in 10X Visium data which is divided by 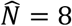.

Cell2location estimates described in this section were reported in Fig 2D-G, Fig 3C-E, Fig S6C-E, Fig S7, Fig S8, Fig S10B-C, Fig S12A, Fig S13, Suppl. Files 1-6.

#### Region identification

Estimated absolute cell abundance was used to calculate a KNN graph (N neighbours=38) followed by Leiden clustering of locations (resolution=1.3), yielding 31 clusters. The clusters were annotated by comparing the paired histology images and clusters marking anatomically similar regions were merged, yielding 29 distinct clusters annotated as tissue regions and reported in Fig 3C-D and Fig S7.

### Analysis of Slide-Seq V2 data

Expression signatures of 59 mouse brain reference cell types (Fig 2B) were derived using the negative-binomial regression model as described above. Slide-Seq V2 data of the mouse brain originating from the region containing the hippocampus, parts of the cerebral cortex and thalamus, was downloaded from the data portal for Stickels, Murray et al^25^: https://singlecell.broadinstitute.org/single_cell/study/SCP815. Genes with mRNA count > 0 in at least 500 locations were retained to reduce data size. One Slide-Seq V2 section with the largest number of spatial locations (9,069 genes shared with the scRNA-seq dataset, 53,208 locations) was used as input to cell2location using the following parameters:

- Training iterations: 30000,
- Cells per location 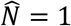, Cell types per location 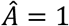, Co-located cell type groups per location 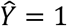, assuming that most locations have only one cell and one cell type on average, accounting for much smaller location size which is similar to the size of single cells (10 microns).
- Prior on the difference in sensitivity between technologies: *μ* = 1/10, *σ*^2^ = 1/400.

Cell2location estimates described in this section were reported in Fig 2H, Fig S10, Suppl. Files 7-8.

### Validation mouse brain cell type locations with region-specific scRNA-seq reference

For quantitatively validating the cell type maps produced by cell2location, we considered a publicly available^24^ gold-standard reference of cortical and hippocampal cell locations. In that data, individual layers of the cortex and the hippocampus are dissected and subjected to scRNA-seq using SmartSeq 4 protocol (Fig 2I). Data from this study was obtained from https://portal.brain-map.org/atlases-and-data/rnaseq/mouse-whole-cortex-and-hippocampus-smart-seq. Cells originating from the primary somatosensory cortex (SSp, n=98) as well as exclusive to the hippocampus (the subiculum areas indicated as PAR-POST-PRE, n=23) were selected, discarding cell types that contribute less than 3 cells. The count of cells in this scRNA-seq data was used to compute the proportion of each reference cell type in each cortical layer. We derived corresponding reference signatures of 121 cell types as per-cluster average expression, as the negative-binomial regression model is not suitable for SmartSeq data (Supp. Methods).

We mapped cell types from this reference to a subset of the mouse brain 10X Visium data, restricted to the primary Somatosensory cortex (which was annotated based on histology). Data was processed as described above and the untransformed mRNA counts (13566 genes shared with scRNA-seq dataset, 1393 locations) were used as input to cell2location using the following parameters (the rest selected as for the primary mouse brain analysis):

- Cell types per location 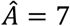;
- Prior on the difference in sensitivity between technologies: *μ* = 1/20, *σ*^2^ = 1/1600, reflecting that the average total mRNA count in a highly sensitive Smart-Seq 4 scRNA-seq is about 20 times greater than the average total mRNA count in 10X Visium data divided by 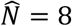.

To quantitatively assess the consistency of the estimated relative cell abundance (Fig 2J, Fig S9C), cell2location estimates of absolute cell density were averaged and normalised per cortical layer to obtain the proportion of cells per cell type within each cortical layer. These estimates were then compared to the cell proportions in the reference scRNA-seq data (Fig 2J), considering 23 most prevalent cell types exclusive to the somatosensory cortex (cell count in SSp > 15, cell count in the hippocampal region = 0 in the scRNA-seq data).

To quantitatively assess the specificity and sensitivity of cell2location at excluding cell types that should be absent in the spatial data, the precision-recall was quantified on all 121 cell types and 5 Visium tissue sections jointly. The 99.5% quantile of mRNA abundance distribution for each cell type and experiment was used as a predictor. The cell count > 0 in the somatosensory cortex region was used as a gold standard label (Fig S9A left panel).

### Identification and spatial mapping of astrocyte subtypes in the mouse brain

To identify regionally enriched astrocytes, we annotated and spatially mapped refined reference signatures of astrocyte subpopulations. Cells annotated as astrocytes were selected from the initial analysis (Fig 2B), as identified based on the expression of canonical markers (e.g. *Aldoc*, *Slc1a3*). These cells were subjected to analysis workflow in scanpy^55^ as follows: First, untransformed count matrix was filtered to select expressed genes at 2 cut-offs:

1. selecting genes detected at mRNA count > 0 in > 5% of cells;
2. selecting genes detected at mRNA count > 0 in a few cells (5% > cell count > 0.4%) but with a mean expression across non-zero cells >1.122018.

The filtered data was then log1p-transformed and scaled. PCA was performed (30 components) and the first component, which was strongly associated with total mRNA count per cell, was removed. Next, BBKNN^33^ was used to refine the integration across all 6 snRNA-seq datasets with the default parameters except, neighbors_within_batch = 3. The corrected KNN graph was used to perform UMAP dimensionality reduction and Leiden clustering with resolution 2.4 identified 17 astrocyte subclusters (Fig S11). Marker genes of these subtypes were identified using differential expression tools in the scanpy package and markers known from the literature.

Next, an initial mapping with cell2location was performed using hyperparameters described above in the mouse brain Visium section. Reference cell type signatures for initial mapping included 17 astrocyte subclusters as well as all other non-astrocyte cell types shown in Fig 2B. Clusters that were not sufficiently distinct in their location (Fig S12) and marker genes (Fig S11) were merged. This resulted in 10 molecularly and spatially distinct astrocyte subtypes (Fig 3A-D). Resulting in 10 merged subtypes were incorporated into cell annotation used throughout the paper. Clustering of locations demonstrated in Fig 3D was performed as described above.

### Analysis of lymph node data

#### Construction of single cell reference data

Published scRNA-seq datasets of lymph nodes have typically lacked an adequate representation of germinal centre-associated immune cell populations due to age of patient donors^56,57^. We, therefore, included scRNA-seq datasets spanning lymph nodes, spleen and tonsils in our single-cell reference to ensure that we captured the full diversity of immune cell states likely to exist in the spatial transcriptomic dataset (Fig 4A-B). The scRNA-seq reference was combined from 3 studies^35–37^. Integration was performed using Scanpy^55^ including removal of doublets with scrublet^53^, regressing out batch from PC space and performing BBKNN. The integrated reference was used to harmonise cell type and subtype labels between datasets, yielding 34 annotated cell populations represented across 73,260 cells.

#### Spatial data

10X Visium spatial transcriptomics data of human lymph nodes was downloaded from the 10X Genomics website: https://support.10xgenomics.com/spatial-gene-expression/datasets/1.0.0/V1_Human_Lymph_Node.

#### Spatial mapping using cell2location

Expression signatures of 34 cell populations (Fig 4B) were derived using the negative binomial regression model (Suppl. Methods) accounting for the effect of different batches as well as technologies (Fig S17). Prior to model-based estimation, untransformed count matrix was filtered to select expressed genes at 2 cut-offs:

1. selecting genes detected at mRNA count > 0 in > 3% of cells;
2. selecting genes detected at mRNA count > 0 in a few cells (3% > cell count > 0.1%) but with a mean expression across non-zero cells > 1.122018.

The untransformed mRNA counts from a single Visium experiment (10,241 genes shared with the scRNA-seq dataset, 4039 locations) were used as input to cell2location using the following parameters:

- Training iterations: 30000,
- Cells per location 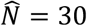, derived by examining histology images of lymph node tissue (paired image had insufficient image quality), Cell types per location 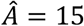, representing a high degree of spatial interlacing of cell type locations in this tissue, Co-located cell type groups per location 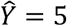, representing a high degree of linear dependencies - that many cell types co-locate and form tissue zones.
- Prior on the difference in sensitivity between technologies: *μ* = 1/3, *σ*^2^ = 1/36.

#### NMF

Cell abundance estimated by cell2location were used as input for non-negative matrix factorization to identify spatially interlaced tissue zones/compartments of lymph nodes (Fig 4D). Cell abundance estimates *w_s,f_* are modelled as an additive decomposition:

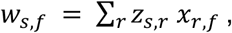

where *x_r,f_* represents NMF weight describing the contribution of each cell type group *r* to the abundance of cell types *f*; *z*_s,r_ represents NMF weight describing the abundances of each cell type group *r*across locations *s*.

NMF was trained for a range of *R* = {8, . ., 28} and the decomposition into 14 factors was chosen as a balance between capturing fine tissue zones and splitting known compartments such as Light Zone of the Germinal Center into several distinct factors (Fig 4E, panel 3) into separate zones. NMF weight *x*_r,f_ is shown in Fig 4D and Fig S16C.

Cell2location estimates from this section are reported in Fig 4, Fig S10B, Fig S16.

### Single-molecule fluorescent in situ hybridization

A wild-type adult C57BL/6 mouse (postnatal day 63, male) was transcardially perfused with ice-cold phosphate-buffered saline (PBS) and 4% paraformaldehyde (PFA) in 1X PBS. The brain was dissected and post-fixed in 4% PFA at 4°C overnight and cryoprotected in 30% sucrose for 48 h at 4°C. The brain was then embedded in OCT (Tissue-Tek) and frozen on isopentane-dry ice slurry and stored at −80°C. Coronal cryosections (12 *μ*m) were collected on a cryostat (Leica), mounted onto superfrost glass slides (VWR) and stored at −80°C.

RNAScope smFISH was performed as previously described^58^. Tissue slides were thawed at room temperature for 15 min and baked at 60°C for 30 min. Afterwards, slides were post-fixed for 15 min in chilled 4% PFA, and serially dehydrated through 50%, 70%, 100%, and 100% ethanol for 5 minutes each. 4-plex RNAScope smFISH was automated using a Leica BOND RX (LS Multiplex Assay, Advanced Cell Diagnostics (ACD), Bio-Techne). Tissue sections were permeabilized using heat (5 minutes in Leica BOND ER2 buffer at 95°C) and protease digestion (ACD Protease III for 20 minutes). The probes in C1, C2 and C3 channels were labeled using Opal 520, 570 and 650 fluorophores (Perkin Elmer, diluted 1:1500) respectively. The C4 probe was developed using TSA biotin (TSA Plus Biotin Kit, Perkin Elmer, 1:500) and streptavidin-conjugated Atto-425 (Sigma Aldrich, 1:400). Nuclei staining was performed with DAPI (Life Technologies Ltd, 1:50,000).

Imaging was performed on an Opera Phenix High-Content Screening System (Perkin Elmer) in spinning disk confocal mode using a 20× water-immersion objective (NA 0.16, 298.99 nm/pixel) with a 1 *μ*m z-step size. The excitation (Ex) laser and emission (Em) filter wavelengths were: DAPI (375 nm; 435-480 nm), Atto-425 (425 nm; 463-501 nm), Opal 520 (488 nm; 500-550 nm), Opal 570 (561 nm; 570-630 nm), Opal 650 (640 nm; 650-760 nm). Illumination correction and stitching were performed on maximum z-projection images using Acapella scripts provided by Perkin Elmer.

### Astrocyte segmentation and smFISH quantification

The segmentation of astrocytes was performed on the *Agt* image channel with pixel classification of Ilastik^59^. The Agt channel was gaussian blurred with a sigma of 2 pixels and the astrocyte mask was subsequently generated by the trained pixel classifier. The performance of this classifier was evaluated by using visual inspection and a metric called out-of-bag(oob) error which measures prediction error by using bootstrap aggregating. The oob error of the classifier was 2.857% measured on a training image of dimension 402 * 2010 pixels. As Agt is expressed at low levels in astrocytes outside the diencephalon^27^, we manually traced the diencephalon to filter all astrocyte signals outside.

Two rounds of “closing” operations were then executed to connect as many major astrocyte cellular processes to the cell bodies as possible. As a consequence, some cells were erroneously fused together. To split these fused cells, we performed six rounds of opening on large cells (>10,000 pixel^2^). Finally, all small partially segmented cells (<250 pixel^2^) were filtered out and the mean pixel intensities (sum intensity/area) of target gene channels were quantified per cell.

## Data availability

snRNAseq data are publicly available via cellxgene portals^60^ at https://cell2location.cellgeni.sanger.ac.uk/mouse-brain (full dataset, annotation_1_print column denotes cell types) and https://cell2location.cellgeni.sanger.ac.uk/mouse-brain-astrocytes (astrocyte subclusters). snRNA-seq and Visium data of the mouse brain as well as the integrated secondary lymphoid organ scRNA-seq data are publicly available for download through S3 buckets at https://cell2location.cog.sanger.ac.uk/browser.html (mouse brain -/tutorial/; lymph nodes-/paper/integrated_lymphoid_organ_scrna/).

## Code availability

The cell2location package is available at https://github.com/BayraktarLab/cell2location/. Documentation and tutorials are available at https://cell2location.readthedocs.io/. Singularity environment for cell2location is available from https://cell2location.cog.sanger.ac.uk/browser.html?shared=singularity/. Code used to generate synthetic data is available in https://github.com/emdann/ST_simulation/. Code used to segment nuclei in histology images is available in https://github.com/yozhikoff/segmentation. Jupyter notebooks covering the analysis in this paper will be made available at https://github.com/vitkl/cell2location_paper/ and upon request.

## Acknowledgements

We thank Britta Velten, Yuanhua Huang and Luca Marconato for feedback on the cell2location model, Martin Prete and Vladimir Kiselev for dockerising the cell2location tool and creating the web portals for sharing our data, Natsuhiko Kumasaka for helpful comments on single cell analysis, Stephen Leonard and Krzysztof Polanski for help with spatial and single nucleus data processing, Kylie James and Rasa Elmentaite for help preparing lymph node single cell datasets, Kenny Roberts for advice on smFISH, Jing Eugene Kwa for advice on snRNA-seq, Jana Eliasova for illustrations and logo design, and David Rowitch and Sarah Teichmann for comments on the manuscript. H.W.K. was funded by a Sir Henry Wellcome PostDoctoral Fellowship (213555/Z/18/Z). This study was supported by Wellcome Trust Core Funding to O.A.B..

## Author Contributions

V.K., O.S. and O.A.B. conceived the study. V.K. developed the cell2location model with feedback from O.S. and worked on validation, mouse brain and lymph node analysis. E.D. worked on generating synthetic data, mouse brain cell annotation and validation of cell type mapping, and contributed to lymph analysis. A.S. implemented cell2location training in PyTorch and pyro, performed Slide-Seq analysis, wrote data visualisation modules and contributed to mouse brain analysis. A.S. and V.KE. contributed to the development and interpretation of the model for estimating cell types signatures. A.L., A.AI. and M.S.J. contributed to the design and interpretation of the cell2location model as well as downstream clustering and NMF. A.AR. performed benchmarking of stereoscope, supervised by R.V.T. and O.S.. H.W.K. contributed to lymph node data analysis and interpretation, supervised by L.J.. L.R. performed snRNA-seq experiments. L.T. performed Visium experiments. J.S.P. performed smFISH experiments and imaging. T.L. performed image processing and astrocyte segmentation analysis. V.K., O.S. and O.A.B. wrote the manuscript with feedback from all authors.

## Supplementary data

**Supplementary figures S1 to S17**.

**Supplementary files 1-5** show spatial cell abundance (colour) maps of 59 cell types (panels) estimated by cell2location across 5 mouse brain tissue sections in 10X Visium data.

**Supplementary file 6** shows spatial mRNA abundance (colour) maps of 59 cell types (panels) estimated by cell2location in the mouse brain section #1 from mouse #1 in 10X Visium data.

**Supplementary files 7-8** show spatial cell and mRNA abundance (colour) maps, respectively, of 59 cell types (panels) estimated by cell2location in Slide-Seq V2 data of the mouse brain.

**Supplementary files 9-10** show spatial cell and mRNA abundance (colour) maps, respectively, of 34 cell types (panels) estimated by cell2location in the human lymph node tissue section in 10X Visium data.

Supplementary files can be accessed at https://github.com/vitkl/cell2location_paper/tree/master/paper/Supplementary_files

