## Supplementary Figures for "Comprehensive mapping of tissue cell architecture via integrated single cell and spatial transcriptomics"

#### Figure S1

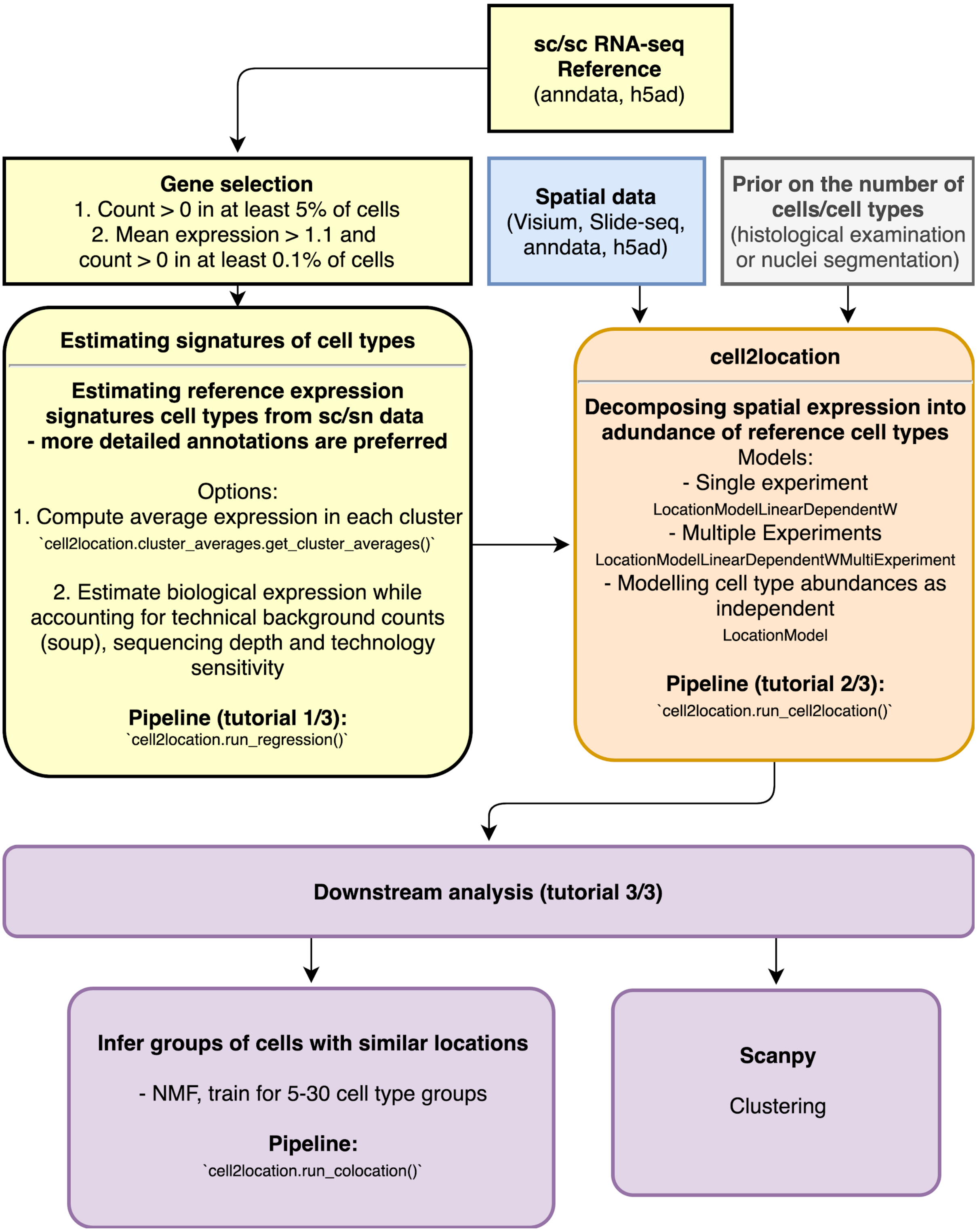


**Figure S1. Details of cell2location workflow.** Diagram of the cell2location workflow, which consists of signature estimation from scRNA-seq data, cell2location cell type mapping and downstream analyses. Each block highlights relevant models, workflows and tutorials.

### Figure S2

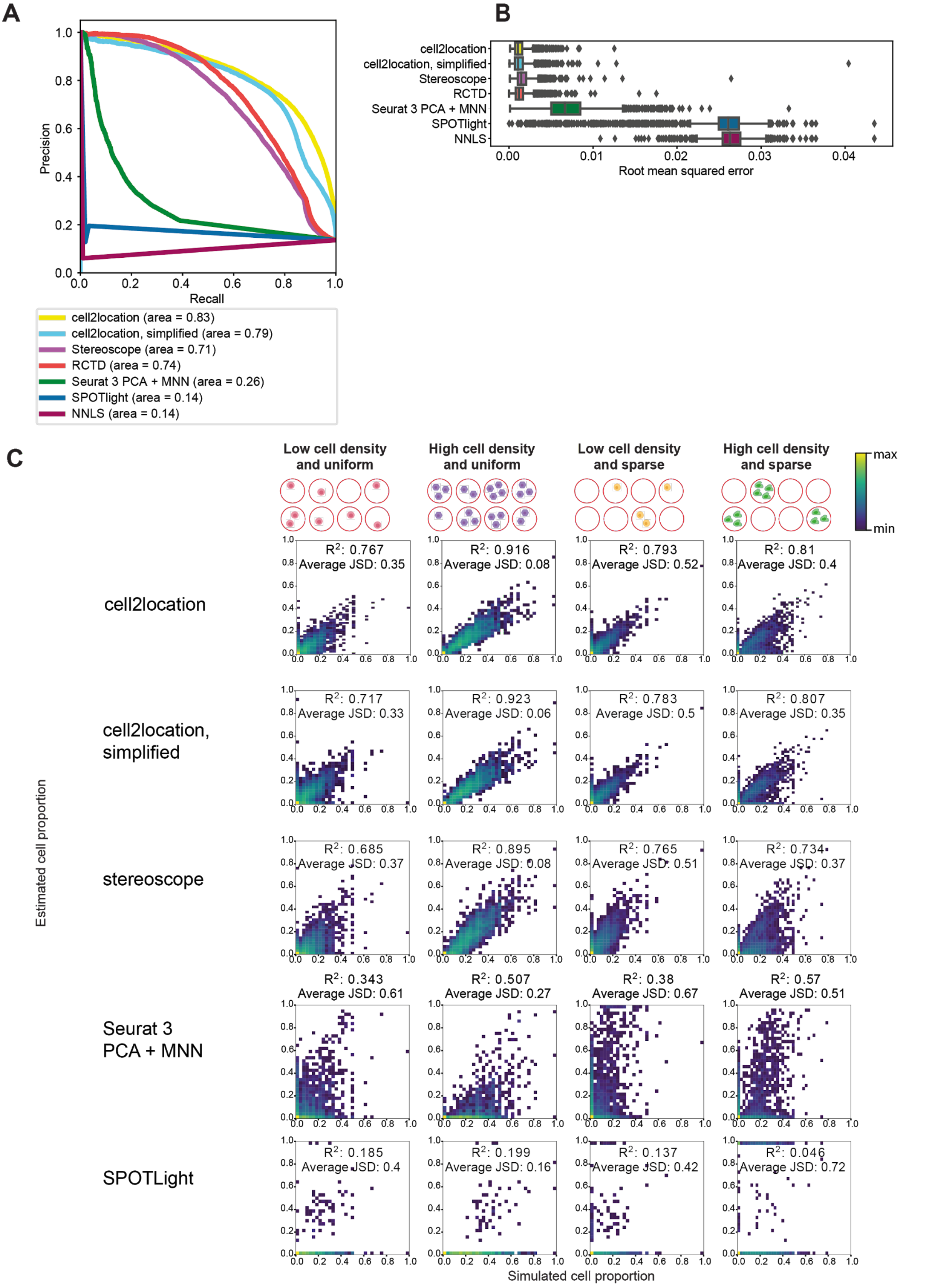


**Figure S2. Assessing the accuracy of cell2location compared to alternative methods.** Assessment of cell2location and alternative models using simulated data. Analogous to the results described in Fig 1. Briefly, the considered benchmark dataset is constructed by combining cells from 46 reference cell types (obtained from mouse brain snRNA-seq, Fig 2B) according to a synthetically generated cell type abundance map.

A. Assessment of cell2location and alternative methods for detecting locations with non-zero cell abundance across all four cell types abundance maps (2,000 locations, 46 cell types). Shown are precision-recall curves for considered methods with the corresponding areas under the curve stated in the legend. Cell2location model accounts for linear dependencies in the abundance of cell types (here termed ‘cell2location’) and showed high accuracy. For reference, we also considered a simpler model, that describes the abundance of cell types as independent, which had lower accuracy for low abundance cell types, confirming the expected benefits of this additional decomposition (Suppl. Methods). Stereoscope was used with default parameters, including selecting 5000 highly variable genes (HVG). RCTD was used with default parameters including selecting cell type marker genes. Seurat V3 was used with default parameters including using PCA rather than CCA for integration as recommended by the authors.
B. Assessment of cell2location and alternative methods at quantitative accuracy of estimated relative cell abundances as measured by root mean squared error (RMSE) across all four cell types abundance maps (2,000 locations, 46 cell types). Boxplot (25%, 75% quantiles, median) shows RMSE (X-axis) for cell2location and alternative methods (Y-axis), the whiskers correspond to the edge of 1.5 inter-quantile-range (IQR) of the lower and upper quartile, separate dots show outliers.
C. Comparison of cell2location and alternative methods across alternative synthetically generated cell type abundance maps. *Top*: alternative cell type abundance maps for data simulation. *Bottom*: 2D histogram plots, displaying the concordance between simulated (X-axis) and estimated (Y-Axis) cell-type proportions across 2,000 locations and cell types (considering 4 cell type abundance patterns; Methods). Each method is shown in rows. Colour denotes 2D histogram counts. $R^2$ denotes Pearson correlation and JSD denotes Jensen–Shannon divergence.

### Figure S3

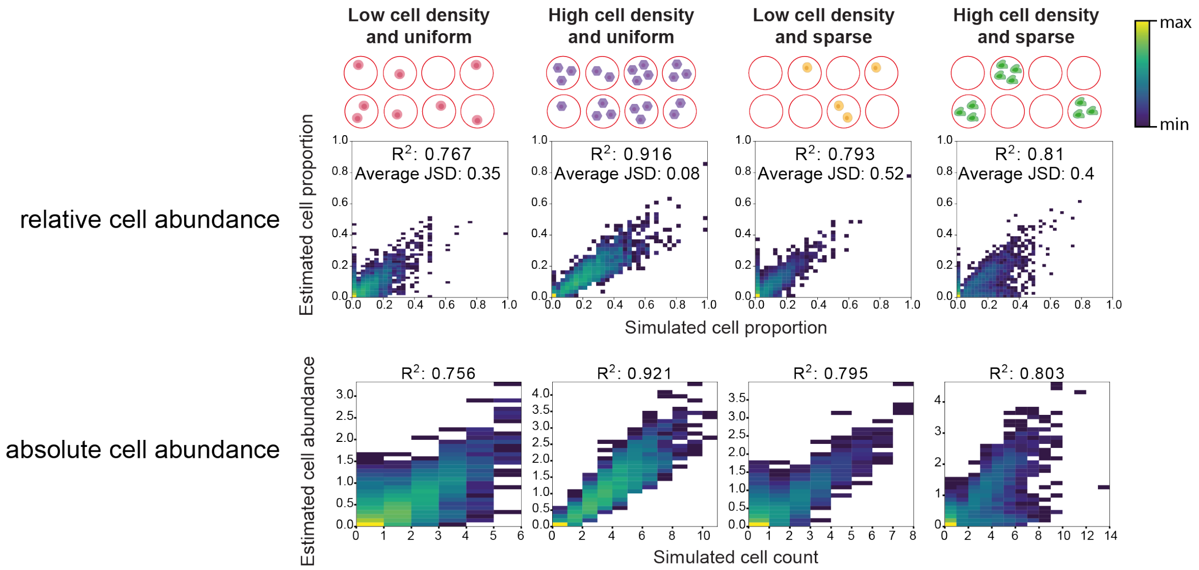


**Figure S3. Assessment of cell2location absolute and relative abundance estimates.** Assessment of the model at estimating absolute cell abundance using simulated data (extending the Fig 1E). Briefly, the considered benchmark dataset is constructed by combining cells from 46 reference cell types (obtained from mouse brain scRNA-seq, Fig 2B) according to a synthetically generated cell type abundance map (Methods).

*Top*: 2D histogram plots, displaying the concordance between simulated (X-axis) and estimated (Y-Axis) cell-type proportions across 2,000 locations and cell types. Colour denotes 2D histogram counts (50 bins along both X- and Y-axis).

*Bottom*: 2D histogram plots, displaying the concordance between simulated (X-axis) and estimated (Y-Axis) absolute cell-type abundance across 2,000 locations and cell types. Colour denotes 2D histogram counts. Each bin along the X-axis corresponds to a discrete cell count in the simulated data (50 bins along Y-axis).

### Figure S4

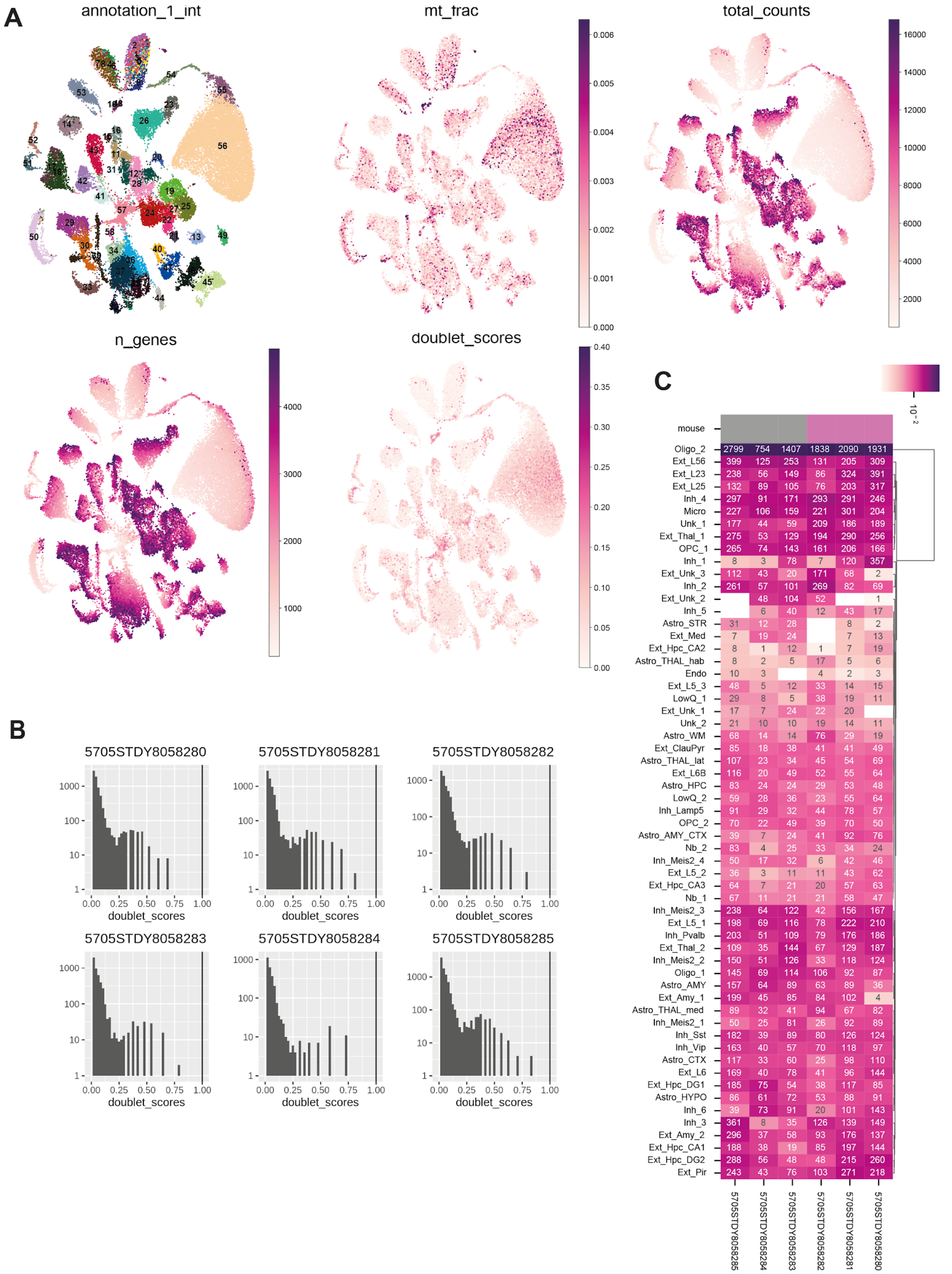


**Figure S4. Quality control metrics of the 59 mouse brain cell type clusters.**

A. UMAP representation (X- and Y-axis) of 59 cell subtypes identified by Louvain clustering on the integrated snRNA-seq dataset across sections (extending Fig 2B). The remaining plots show quality control metrics: the proportion of mRNA counts per cell coming from mitochondrial-encoded genes, total mRNA counts per cell, the total number of genes with mRNA counts > 0, doublet cell scores as identified by scrublet tool.

B. Histogram of doublet scores (X-axis) across 6 profiled mouse brain tissue sections (panels), high values representing a higher likelihood that cell barcode corresponds to a doublet (a droplet containing two cells and one bead with a single cell barcode). The Y-axis of histogram counts is shown on a log scale. Cells with doublet scores > 0.2 were removed from the analysis in panel A and the rest of the paper.
C. Heatmap denoting the proportion of cells (colour, log scale, normalised per tissue section) of each of the 59 cell subtypes (columns) coming from each tissue section (rows). Colour bar denotes mouse 1 (purple) and mouse 2 (grey). Text denotes the count of cells.

### Figure S5

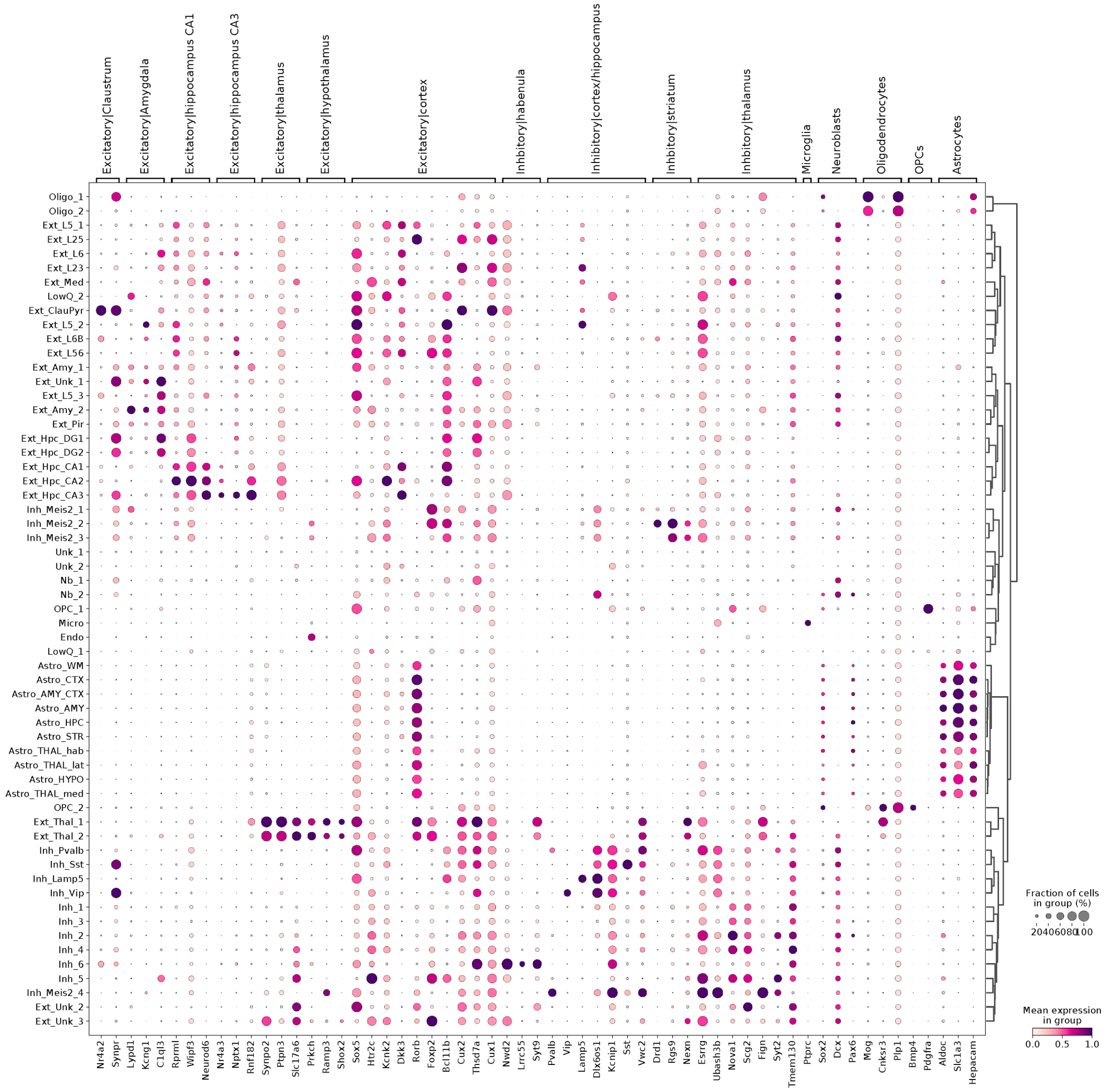


**Figure S5. Regional marker genes of 59 cell type clusters in the mouse brain.** The expression level of regional marker genes for 59 mouse brain cell clusters, including 43 annotated putative cell types and 10 astrocyte subtypes. Dot size denotes the fraction of cells per cluster (rows) that express a given marker gene (count > 0; columns). Dot colour denotes the relative expression of each gene in each cluster, normalised by the maximum values for each gene (scaled between 0 and 1). Marker genes are grouped by region and broad cell type (regional excitatory and inhibitory neurons from Zeisel et al, 2015, 2018 and Tasic et al 2018, as well as glial cell types).

#### Figure S6

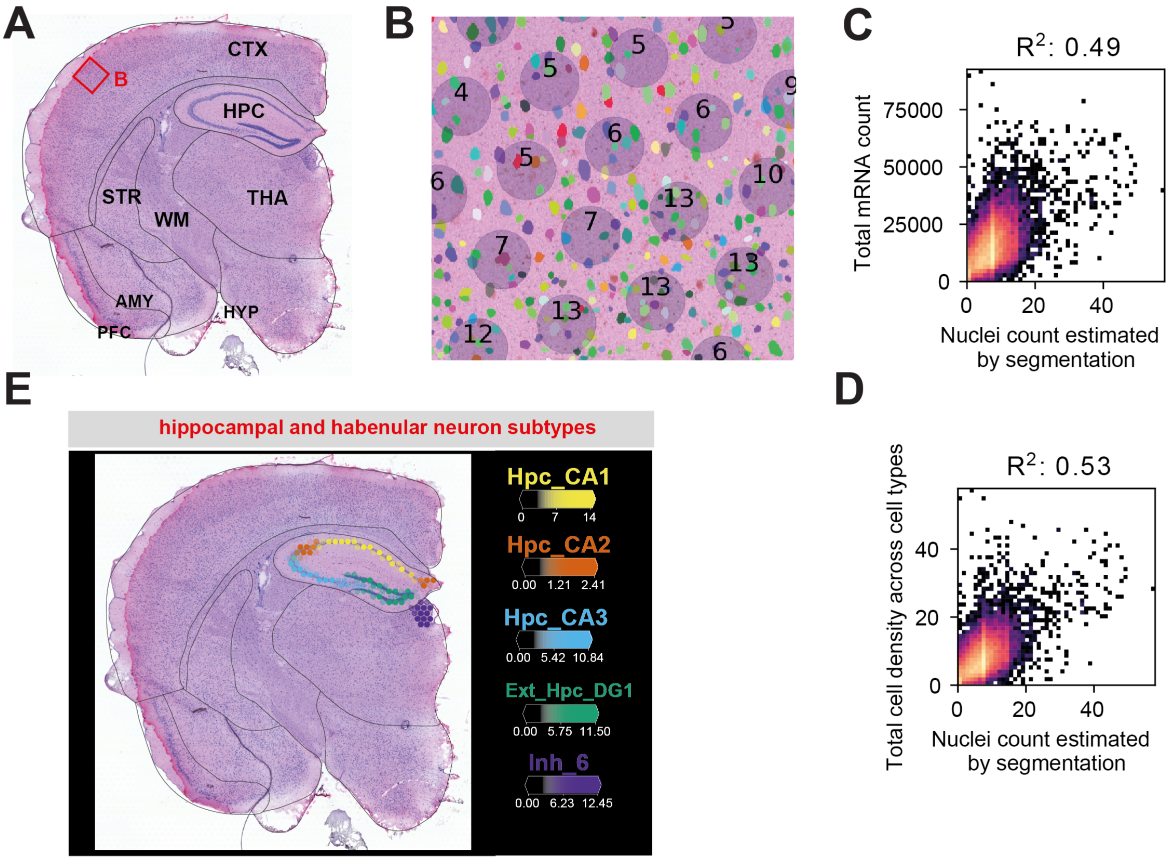


**Figure S6. Cell abundance estimates from image segmentation and cell2location.**

A. Mouse brain regions present in the assayed section. Shown is a histology (H&E) image of the tissue section and the outlines of major brain regions. The red bounding box corresponds to the region in B.
B. A close-up image of the somatosensory cortex. Nuclei segmentation masks (pseudocolours) are shown over H&E image of the tissue section in the background. Grey circles denote the positions and size of the 10X Visium array “spots” (i.e. mRNA capture locations). The numbers denote the number of nuclei that overlap with the capture locations.
C. 2D histogram plots, displaying the correlation between the number of nuclei estimated via segmentation of the H&E image (X-axis) and the total mRNA count per location detected using the Visium technology (Y-Axis). $R^2$ denotes Pearson correlation.
D. 2D histogram plots, displaying the correlation between the number of nuclei estimated via segmentation of the H&E image (X-axis) and the total cell abundance estimated by cell2location (Y-Axis). $R^2$ denotes Pearson correlation.
E. Estimated cell densities (colour intensity) of excitatory neurons across hippocampal divisions (Ext Hpc CA1, Ext Hpc CA2, Ext Hpc CA3, Ext Hpc DG1) and habenular inhibitory subtype Inh 6.

### Figure S7

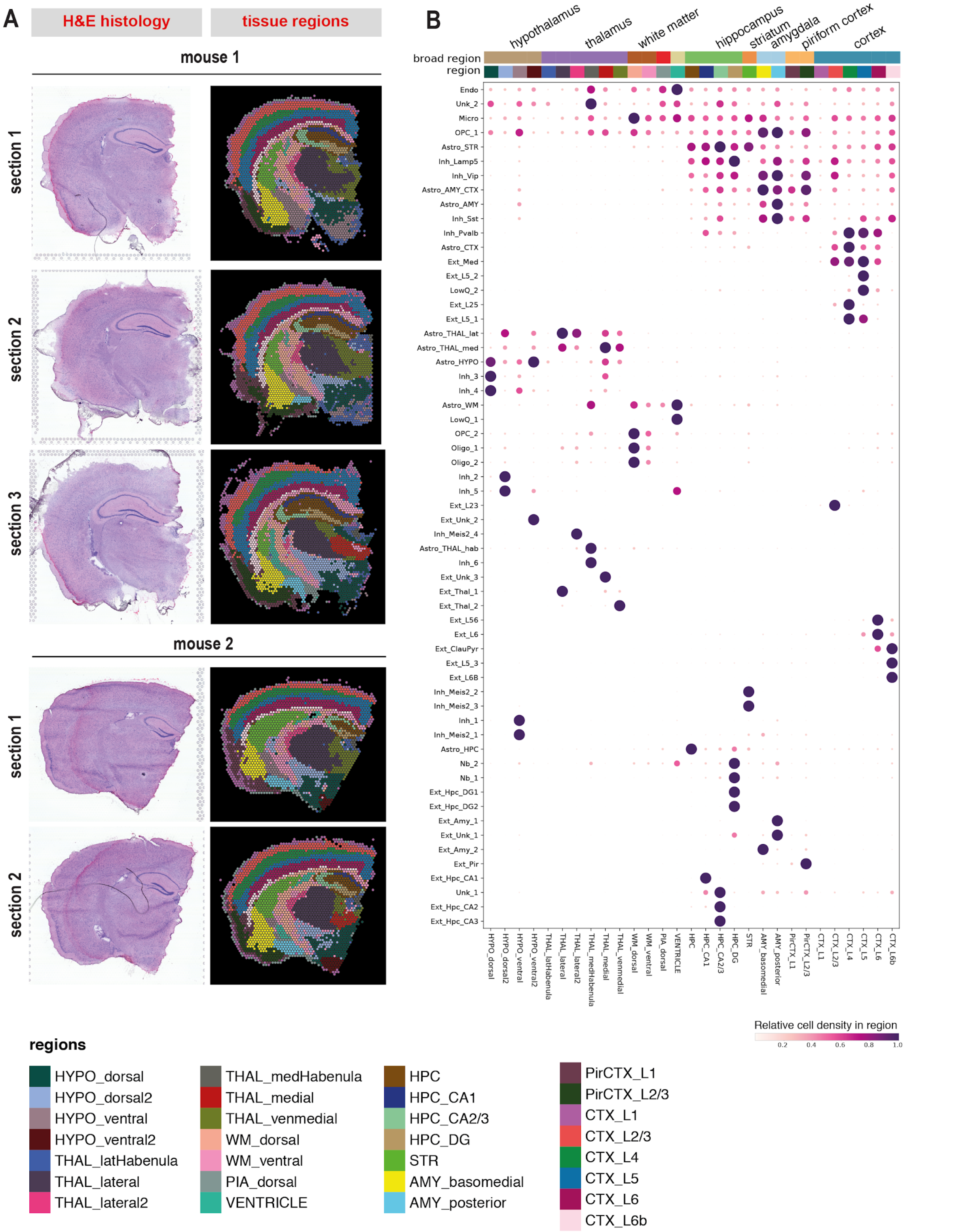


**Figure S7. Distribution of mouse brain cell types across tissue regions identified by clustering.**

A. The distribution of tissue regions identified by clustering locations across adjacent tissue sections. Sections (rows) are grouped by mouse and adjacent sections are approximately 250 microns apart. Shown are histology images (left) and Visium locations and regions identified from clustering (right). The shared legend shows tissue regions identified by clustering (colour). Clustering of locations is based on the cell2location cell type estimates across all slides, followed by anatomical annotation (Methods).

B. Estimated cell densities of 59 cell subtypes across 29 brain regions. Shown is a dot plot, with dot size and colour corresponding to the relative cell abundance for each cell type (rows) across regions (columns), scaled between 0 and 1 by normalising by the maximum values for each cell type. Colour bars (top rows) denote broad brain areas (first row) and subregion (second row, colour code matching panel A).

### Figure S8

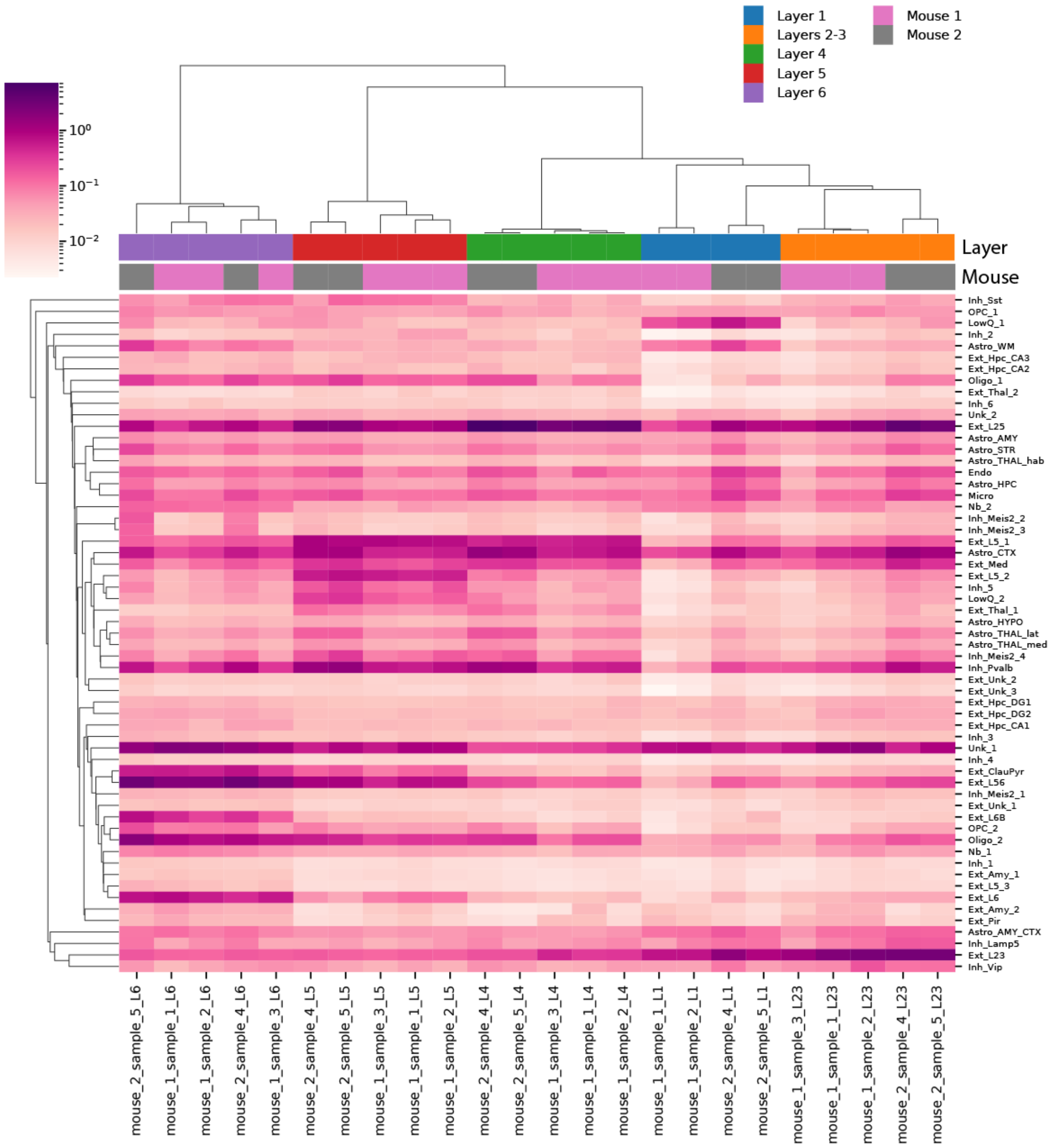


**Figure S8. Hierarchical clustering of cell type abundance estimates recapitulate cortical layers, consecutive sections and replicate mice.** Estimated cell densities of 59 cell subtypes across 5 cortical layers, 3 sections from mouse #1 and 2 sections from mouse #2. The heatmap with a colour corresponding to the absolute cell abundance (log 10 scale) for each cell type (rows) across cortical layers separately in each section (columns). Colour bars denote cortical layers (top) and mice samples were taken from (bottom). The correlation metric and “single” linkage strategy were used for clustering.

#### Figure S9

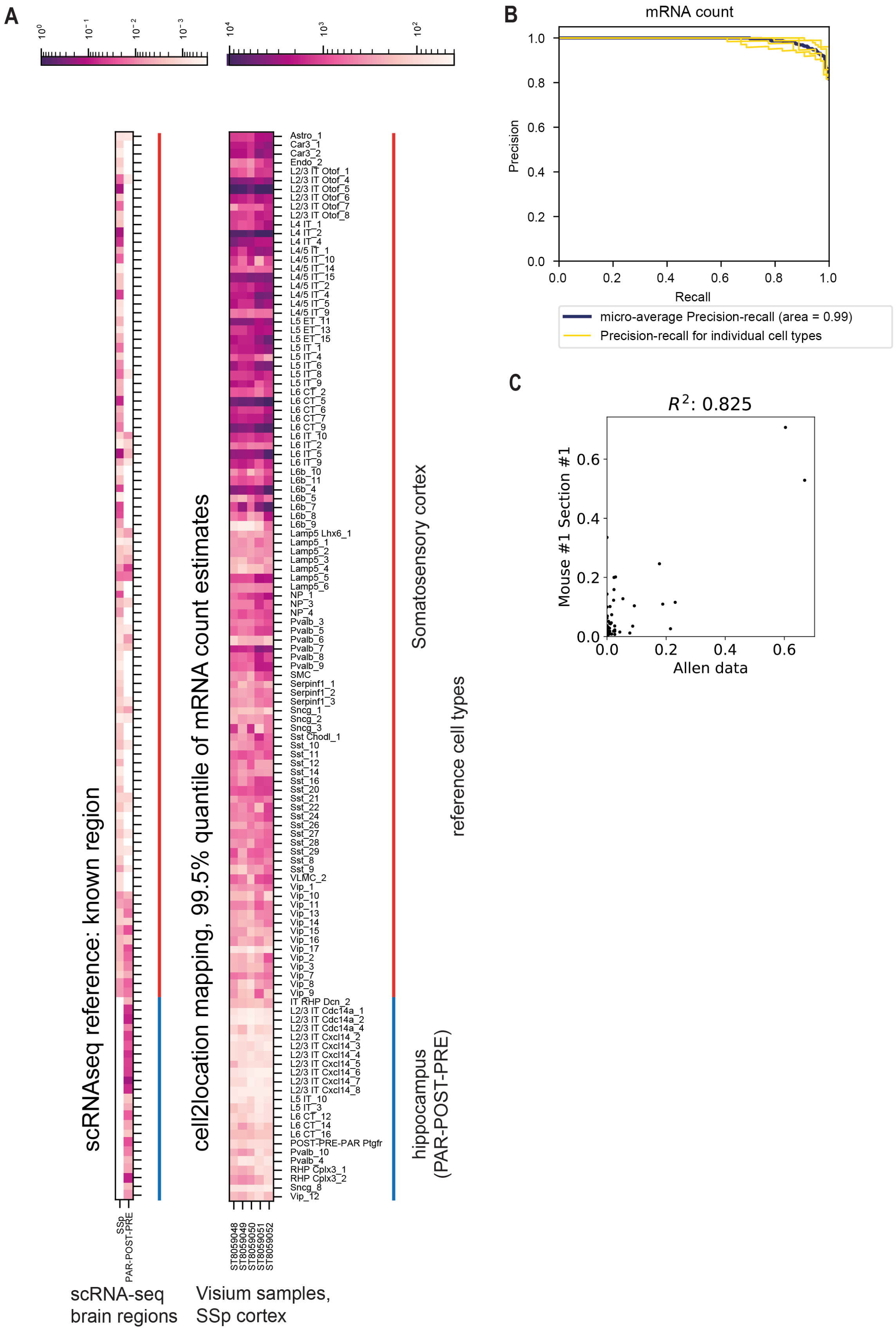


**Figure S9. The accuracy of cell2location at excluding cell types absent in the spatial data (ground truth).** The accuracy of the cell2location at assigning high absolute cell abundance to cell types present in the source brain region (98 subtypes found by Yao et al in somatosensory cortex, SSp) and low cell abundance to cell types absent from SSp region (23 cell subtypes found exclusively in hippocampal brain region by Yao et al).

A. Comparison of cell proportion in scRNA-seq reference (left) and the cell2location mRNA estimates (right). *Left*: Heatmap of relative proportion (colour, log10 scale) of each cell subtype (rows) within the somatosensory cortex (SSp) and selected hippocampal regions (PAR-POST-PRE subiculum) (columns) in scRNA-seq data from Yao et al. *Right*: Heatmap of the 99.5% quantile of mRNA count (colour, log10 scale) from each cell type (rows) in each Visium sample from the primary somatosensory cortex region (columns).
B. Assessment of cell2location at detecting cell types present in the somatosensory cortex region. Shown are precision-recall curves for individual Visium samples (yellow) and their micro-average (blue). 99.5% quantile of mRNA count of each cell type is used as a predictor (panel A right). The cell count > 0 in the SSp region was used as a gold standard label (panel A left). The legend shows the area under the curve.
C. Consistency between relative cell subtype proportion in each cortical layer in the Yao et al reference (X-axis) and the map of those cell types in Visium using cell2location (Y-axis). This scatterplot quantifies the data shown in Fig 2J. Cortical layers and 23 most prevalent exclusive somatosensory cell types are considered; $R^2$ denotes Pearson correlation.

### Figure S10

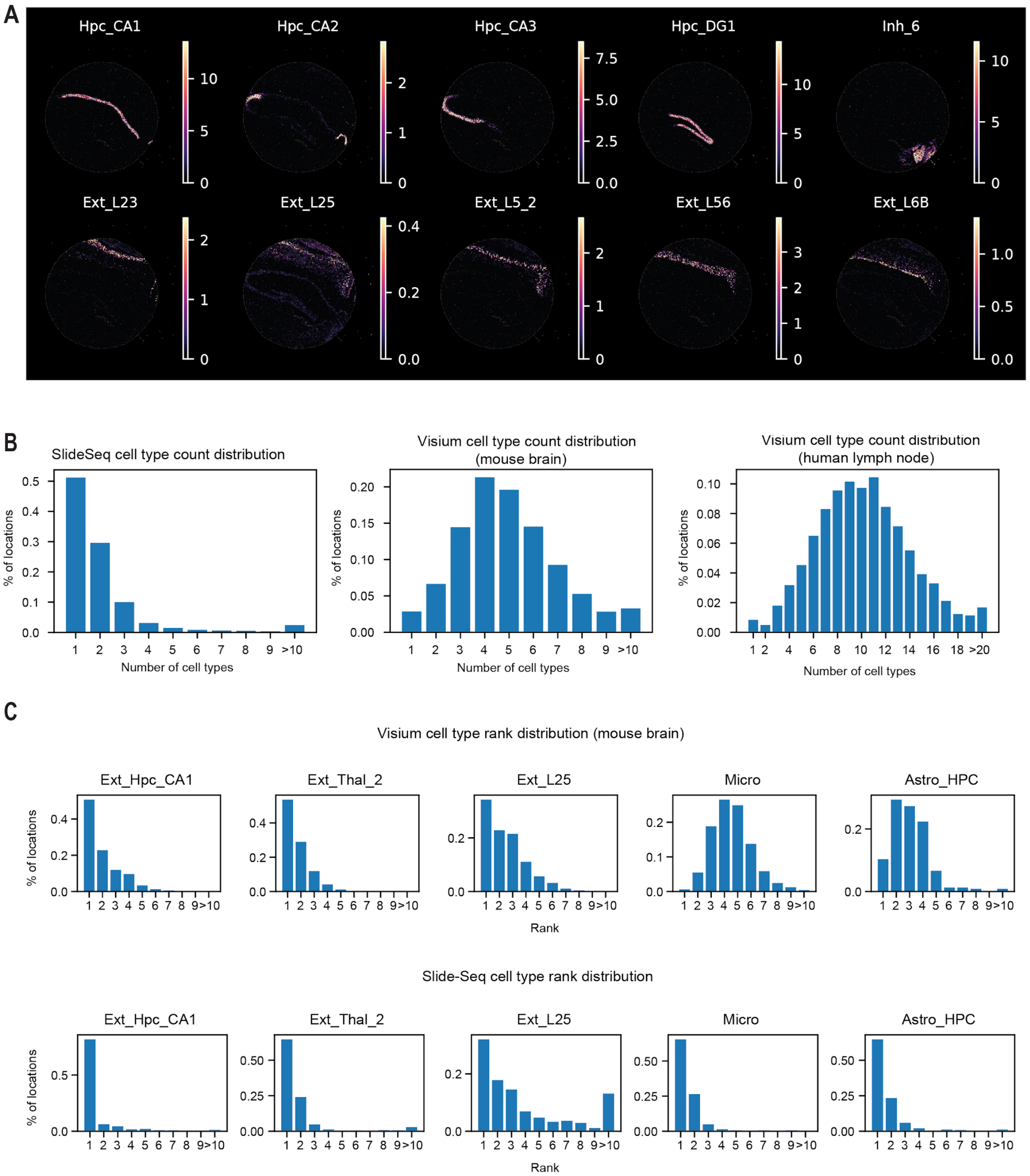


**Figure S10. Comparison of the number of cell types per location in Slide-Seq V2 and 10X Visium data.**

A. Cell2location estimates of cell abundance for 10 cell types (colour) in the hippocampus region (top row) and across cortical layers (bottom row) in the Slide-Seq V2 data (extending Fig2H). Each point represents a Slide-Seq V2 “bead”. Data was clipped to the 99.5% percentile.

B. The histogram (Y-axis) represents the distribution of cell type count (X-axis) across locations (10X Visium “spots” and Slide-Seq V2 “beads”). Cell type is considered to be present in a location if the estimated value of its mRNA count or cell density exceeds a certain threshold which was selected by examining cell type location maps following filtering by this threshold (See below).

*Left*: 59 cell subtypes located in Slide-Seq V2 data of the hippocampus region (Stickels et al), partially covering the cortex and the thalamus, shown in panel A. The threshold is mRNA count > 40. Beads with total mRNA count outside 99.5% quantile were excluded from the analysis as outliers.

*Middle*: 59 cell subtypes located in 10X Visium data of the mouse brain (shown in Fig 2D-F). 4-5 cell types are observed in each location. The threshold is cell density > 0.4.

*Right*: 34 immune cell populations located in the 10X Visium data of human lymph node region (shown in Fig 4). Many cell types (>10) are observed at each location, indicating high spatial interlacing of cell types. The threshold is cell density > 0.8.

C. The distribution of cell type rank in terms of mRNA count across all locations, where the given cell type is present. Cell type is considered to be present if the estimated value of its mRNA count or cell density exceeds a certain threshold. *Top row*: 10X Visium data of the mouse brain, including the hippocampus region, the threshold is cell density 0.5. *Bottom row*: Slide-Seq V2 data of the hippocampus region, the threshold is 60 mRNA. Beads with total mRNA count outside 99.7% quantile were excluded from the analysis as outliers.

### Figure S11

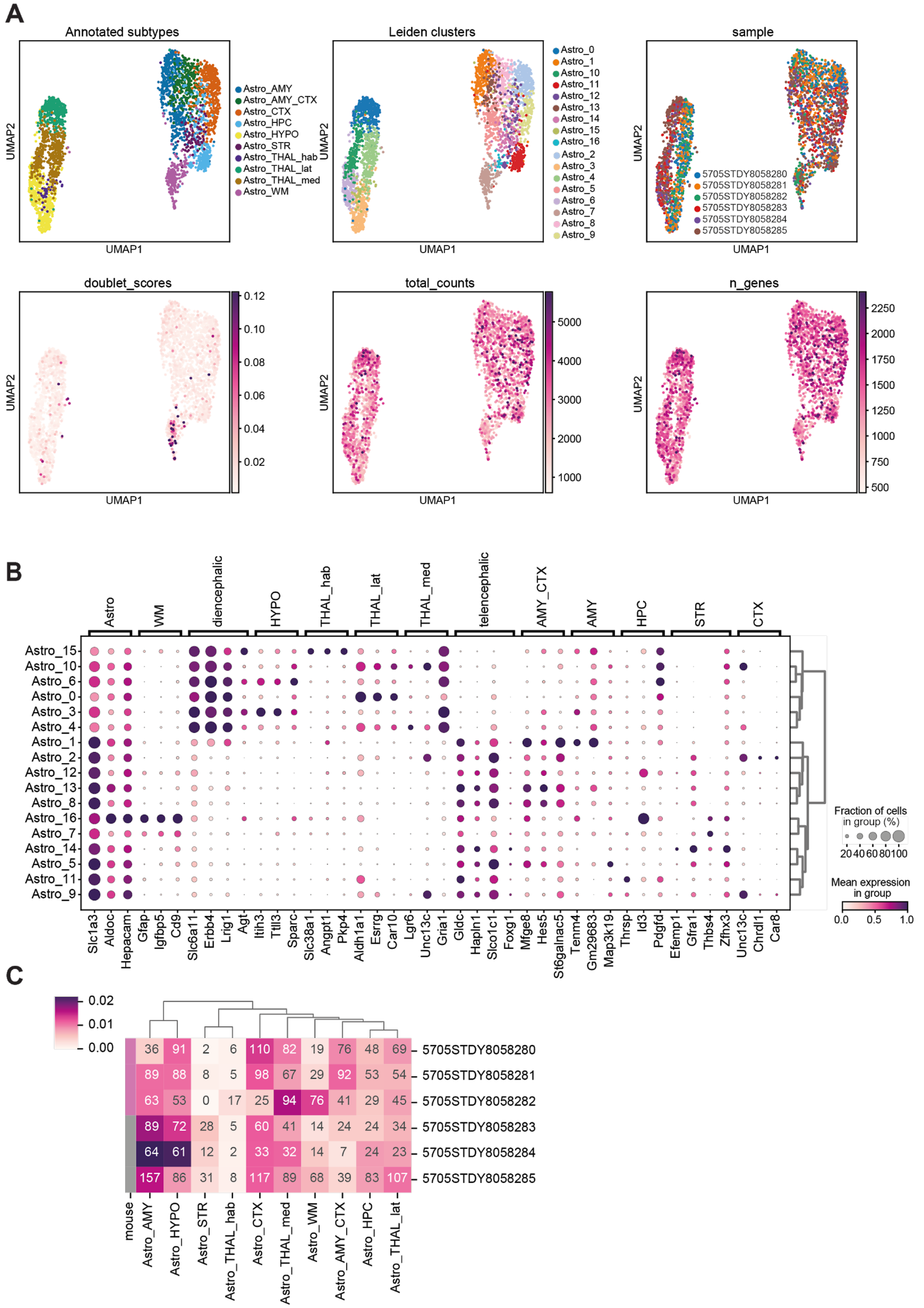


**Figure S11. Mouse astrocyte subtype annotation and quality control.**

A. UMAP representation (X- and Y-axis) of 10 merged astrocyte subtypes compared to the 17 original Leiden clusters (extending Fig 3A). The remaining plots show quality control metrics: doublet cell scores as identified by scrublet tool, total mRNA counts per cell, the total number of genes with mRNA counts > 0.

B. Marker genes of the 17 original astrocyte Leiden subclusters (see their spatial locations in Fig S12). Dot size corresponds to the fraction of cells in each astrocyte subtype cluster (rows) that express a given marker gene (count > 0; columns). Dot colour denotes the relative expression of each gene in each cluster, normalised by the maximum values for each gene (scaled between 0 and 1). Marker genes are grouped by subtypes.

C. Heatmap denoting the proportion of cells (colour, log scale, normalised per tissue section) of each of the 10 merged astrocyte subtypes (columns) coming from each tissue section (rows). Colour bar denotes mouse 1 (purple) and mouse 2 (grey). Text denotes the count of cells.

#### Figure S12

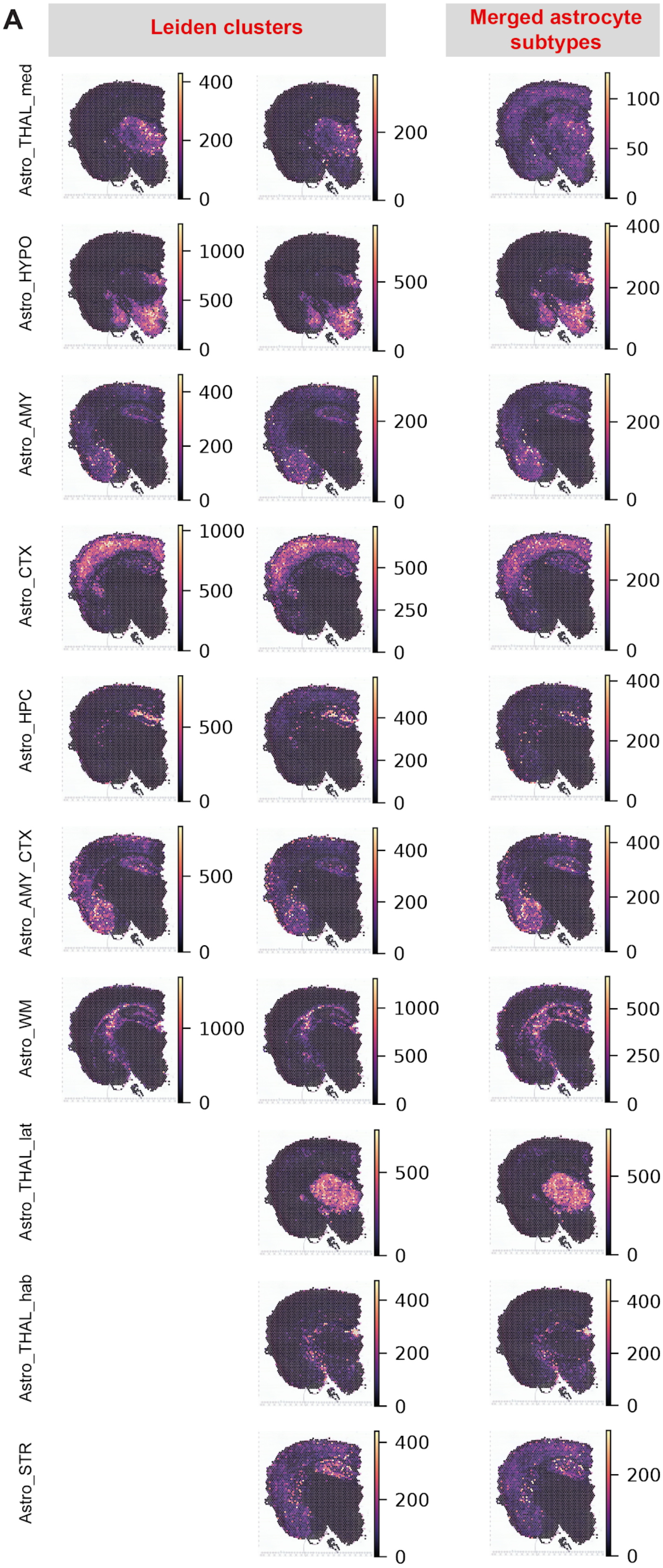


**Figure S12. Refined annotation of astrocyte clusters based on cell2location mapping.** Comparison of absolute mRNA abundance (colour) from each of 10 merged astrocyte subtypes (right, one row per subtype, as reported in Fig 3A) to the cell abundance of original 17 Leiden clusters (left).

#### Figure S13

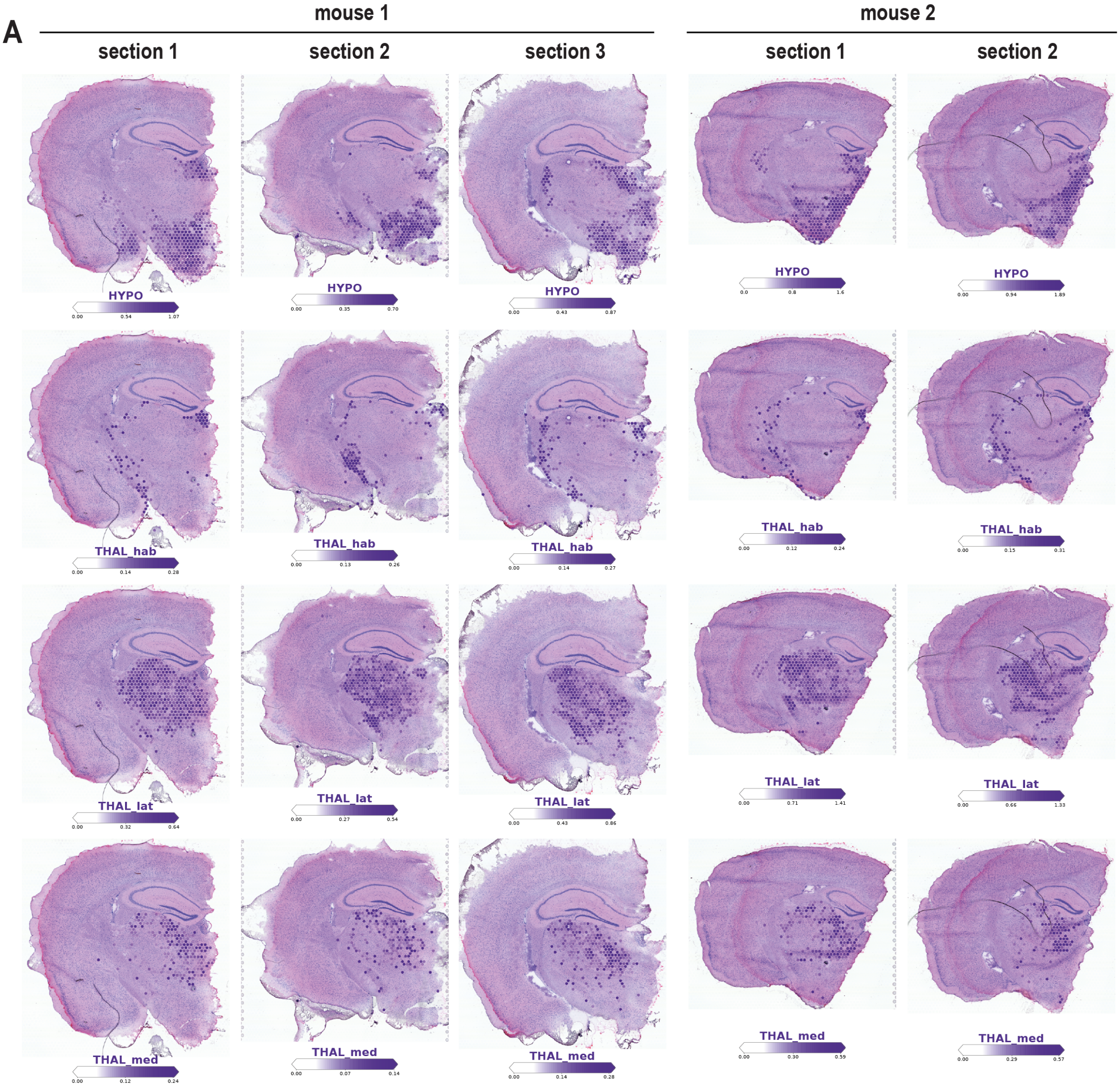


**Figure S13. Estimated locations of diencephalic astrocytes in Visium data.** Estimated cell densities (colour intensity) of 4 diencephalic astrocyte subtypes on tissue sections (approximately 250 microns between consecutive sections). Sections (rows) are grouped by mouse and histology images are shown in the plot background. Rows show HYPO, THAL_hab, THAL_lat and THAL_med astrocyte subtypes, respectively.

### Figure S14

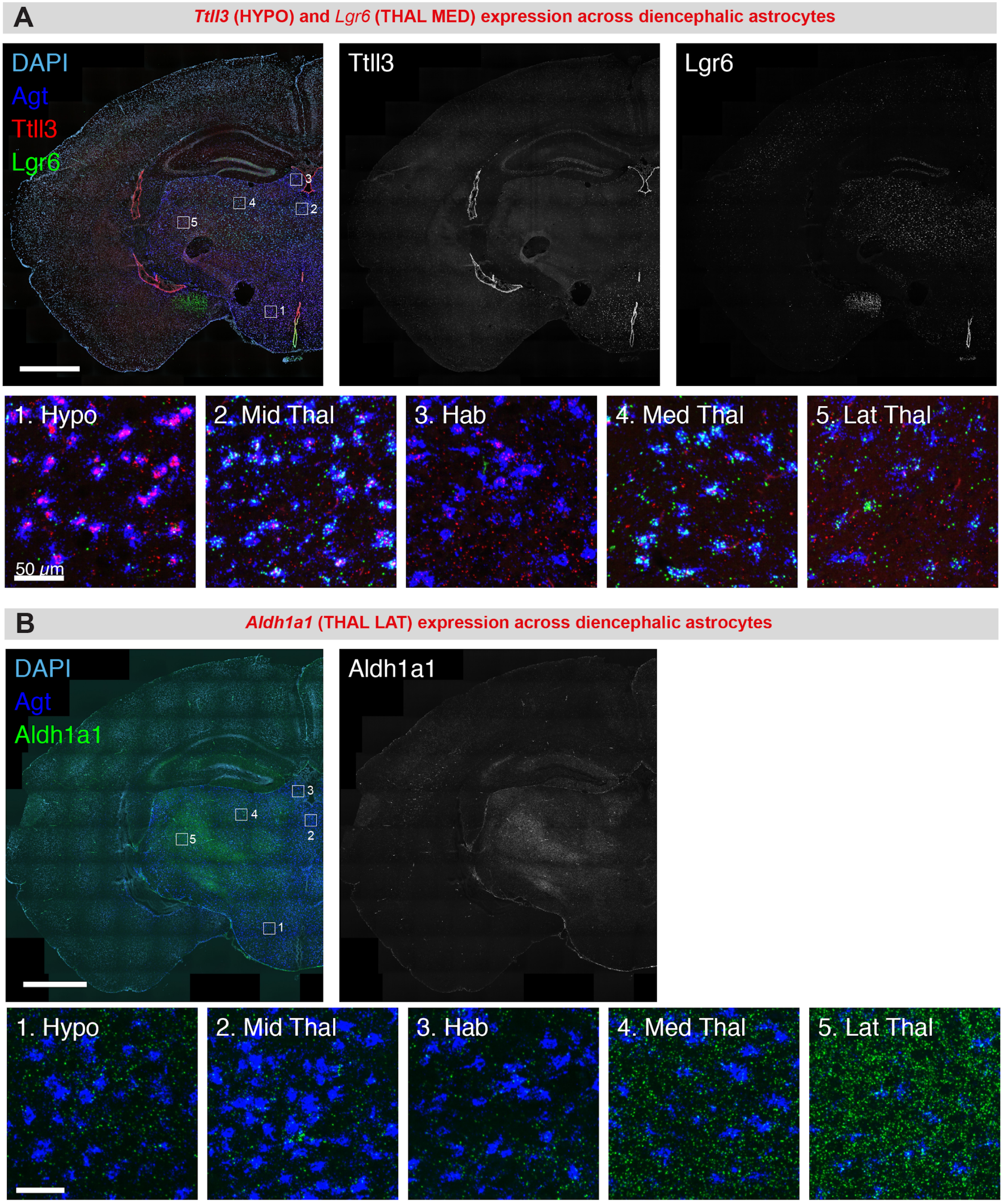


**Figure S14. smFISH validation of hypothalamic and thalamic astrocytes.** RNAScope smFISH validation of regional astrocyte subtype markers across the thalamus and hypothalamus. Large panels show stitched maximum z-projections of coronal hemi-sections. Small panels show close-up images of boxed areas on the large images in the hypothalamus (Hypo), midline thalamic nuclei (Mid Thal), lateral habenula (Hab), medial thalamic nuclei (Med Thal) and lateral thalamic nuclei (Lat Thal). Agt expression marks the soma and major processes of diencephalic astrocytes while DAPI labels all cell nuclei. *Aldh1a1* and *Ttll3* also express in non-astrocyte cell types in the brain. Scale bars: large panels, 1 mm; small panels, 50 microns.

#### Figure S15

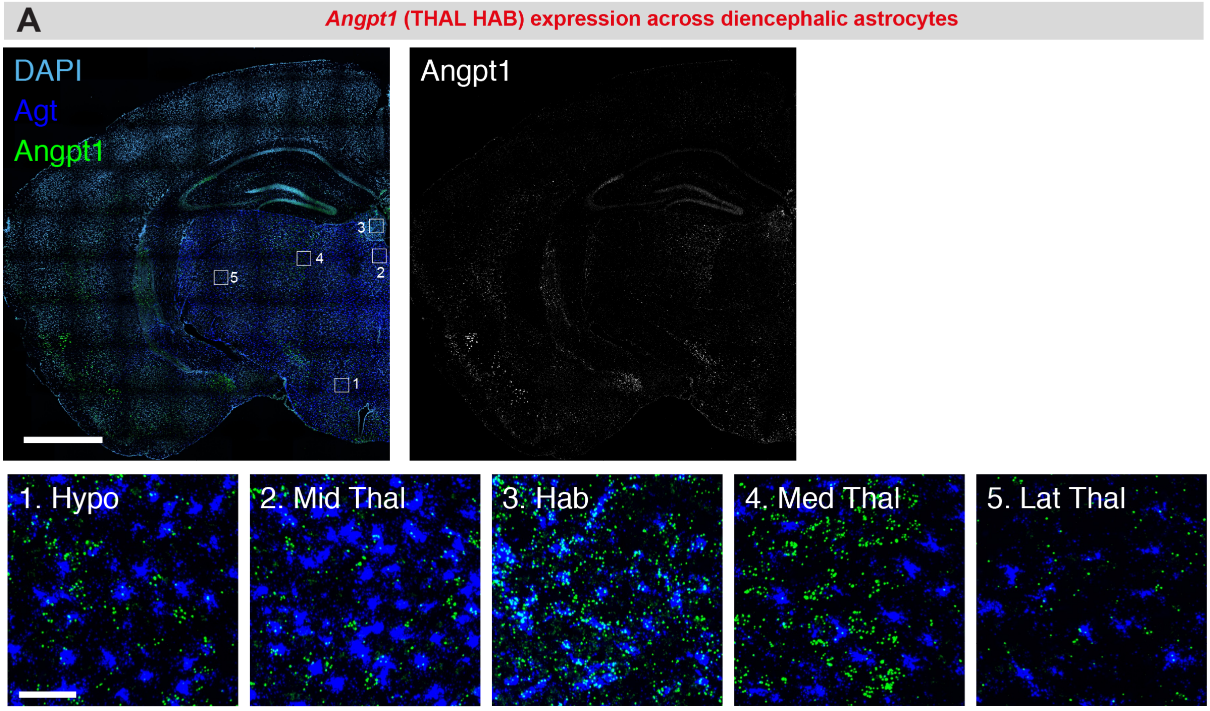


**Figure S15. smFISH validation of habenular astrocytes.** RNAScope smFISH validation of regional astrocyte subtype markers across the thalamus and hypothalamus. Large panels show stitched maximum z-projections of coronal hemi-sections. Small panels show close-up images of boxed areas on the large images in the hypothalamus (Hypo), midline thalamic nuclei (Mid Thal), lateral habenula (Hab), medial thalamic nuclei (Med Thal) and lateral thalamic nuclei (Lat Thal). Agt expression marks the soma and major processes of diencephalic astrocytes while DAPI labels all cell nuclei. *Angpt1* also expresses in non-astrocyte cell types in the brain. Scale bars: large panels, 1 mm; small panels, 50 microns.

### Figure S16

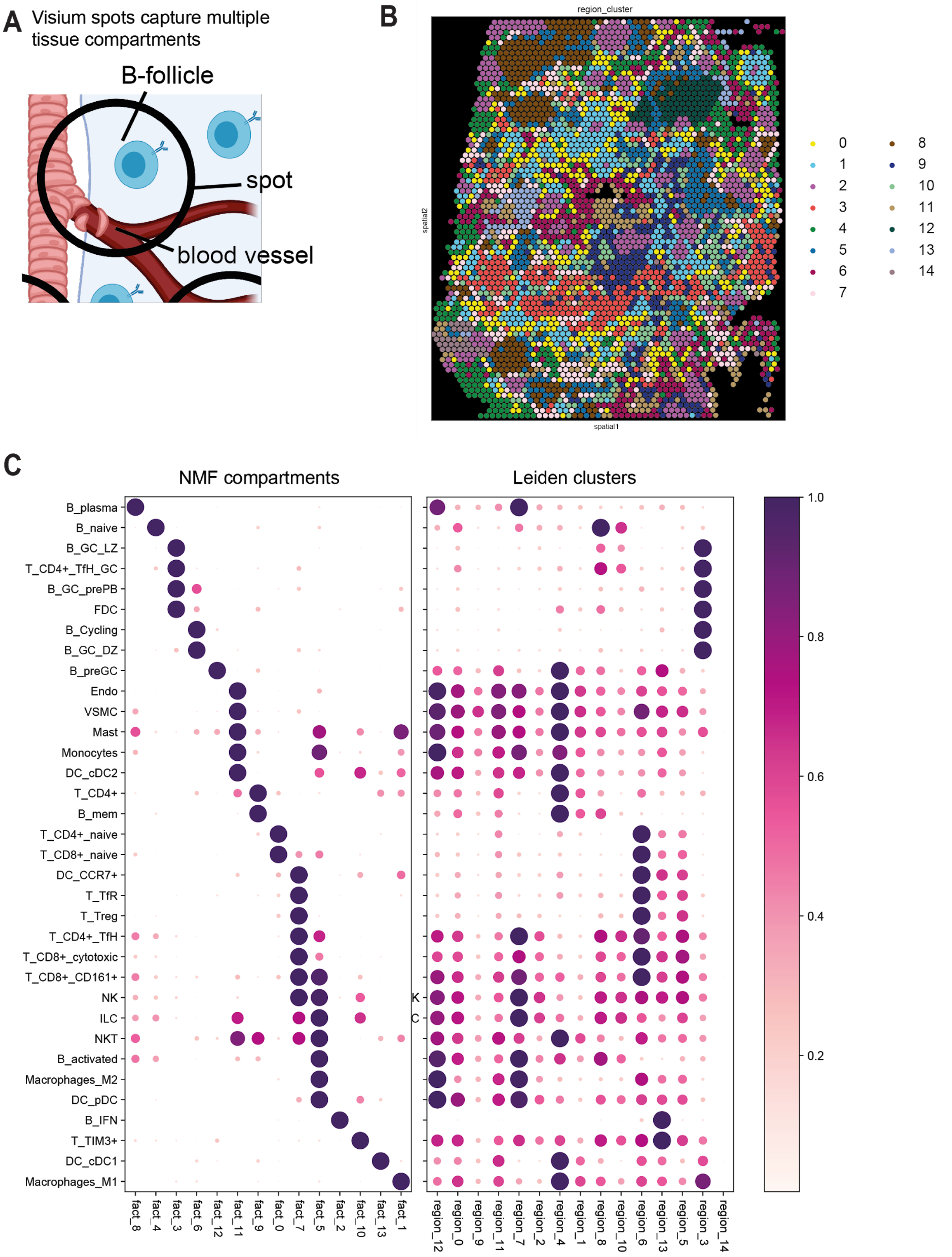


**Figure S16. Comparison of NMF and location clustering for identifying spatially interlaced tissues zones in the lymph nodes.**

A. Diagram illustrating that 10X Visium capture areas are not aligned with tissue compartments and hence tend to capture multiple tissue compartments, such as B-follicle zone and blood vessel zone. This motivates using additive decomposition

(NMF), which naturally accounts for mixed class assignment compared to discrete clustering for identifying such spatially interlaced zones.

B. Human lymph node regions (colour) as identified based on clustering of locations using cell2location cell type abundance estimates, using Leiden clustering with resolution adjusted to produce 14 clusters shown in panel C. Corresponding histology image is shown in Fig 4C.

C. Comparison of cell abundances (rows) across NMF zones (cf. Fig. 4D) and regions identified by Leiden clustering (matching panel B). Shown is a dot plot of the NMF weights (left) for each cell type (rows) in each factor (columns), and average abundance for each cell type in each region(right, columns). Dot size and colour correspond to NMF weights and average abundance in each cluster (normalized across components per cell type and across clusters per cell type).

### Figure S17

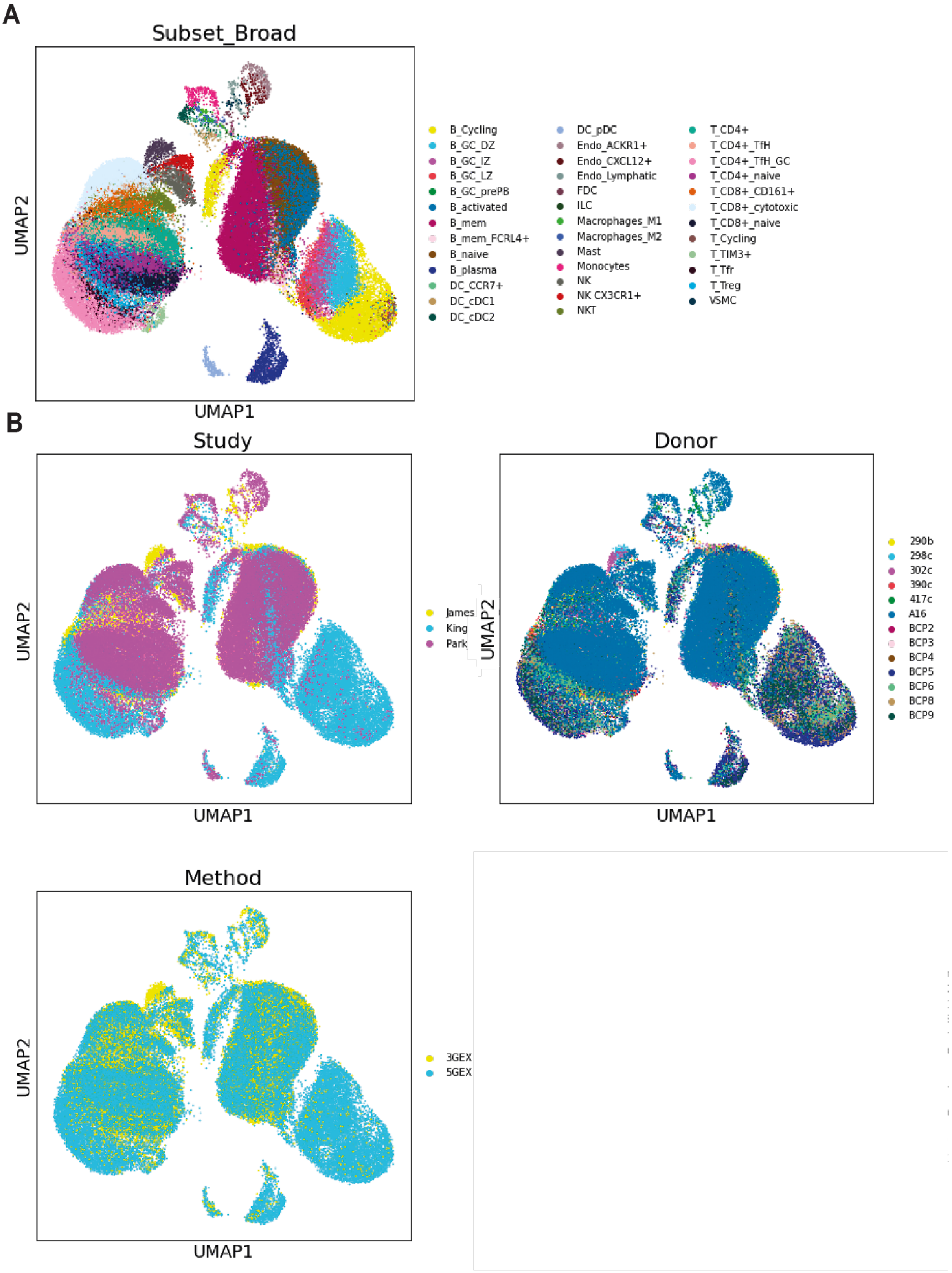


**Figure S17. Construction of reference cell type signatures for the lymph node.**

A. UMAP representation (X- and Y-axis) of 34 immune and non-immune cell subtypes (colour), combining scRNA-seq data from 3 studies (James et al 2020, Park et al 2020, King et al 2020) used to define reference gene expression signatures of cell types.

UMAP was constructed by applying standard Scanpy workflow to scRNA-seq data corrected to remove technology and experiment effect estimated by the regression model (Suppl Methods, section 2).

B. UMAP representation (X- and Y-axis) of the subtypes shown in **panel A**. Each panel shows cells coloured by study (paper reporting the data), the donor, technology (method).
