## Supplementary Methods for "Comprehensive mapping of tissue cell architecture via integrated single cell and spatial transcriptomics"

### Comprehensive mapping of tissue cell architecture via integrated single cell and spatial transcriptomics: cell2location model Supplementary Methods

Vitalii Kleshchevnikov, Artem Shmatko, Emma Dann, Alexander Aivazidis, Hamish W King, Tong Li, Artem Lomakin, Veronika Kedlian, Mika Sarkin Jain, Jun Sung Park, Lauma Ramona, Liz Tuck, Anna Arutyunyan, Roser Vento-Tormo, Moritz Gerstung, Louisa James, Oliver Stegle, Omer Ali Bayraktar

12 November 2020

**2 Estimation of reference expression signatures of cell types** **8**

**3 Comparison of cell2location with alternative methods** **9**

**4 Downstream analyses** **9**

### 1 Mapping cell types with cell2location model

#### 1.1 Cell2location model

Cell2location is a Bayesian model that is aimed at decomposing location-level gene expression counts into a set of pre-defined signatures of reference cell types. A representation as graphical model is shown in Fig 1. The input data of the method are as follows:

Let $D = \{d_{s,g}\}$ be a $S \times G$ untransformed and unnormalised spatial expression count matrix for locations $s = \{1, .., S\}$ and genes $g = \{1, .., G\}$. For 10X Visium data, this matrix can be directly obtained from the 10X SpaceRanger software and imported into data format used in a popular python package Scanpy[1] (see main text Methods). $d_{s,g}$ should be filtered to a set of

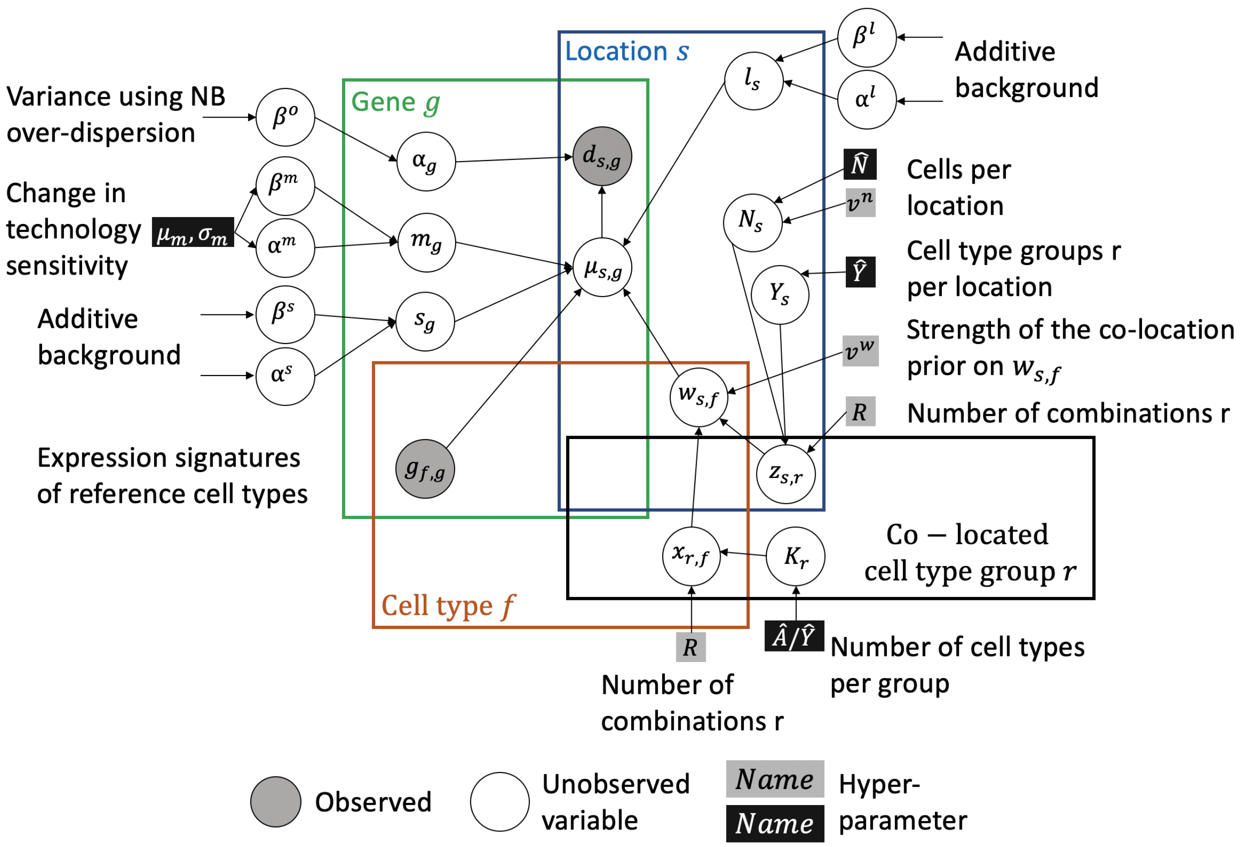

Figure 1: Illustration of the corresponding graphical model. Square boxes denote hyper-parameters, with black box highlighting the parameters that require adaptation to a specific data set (determined by tissue type and spatial technology), as mentioned in Section 1.3.

genes expressed in the single cell reference $g_{f,g}$. The software implementation of cell2location builds on the Scanpy file formats to take advantage of data handling and visualisation tools. For other technologies, such as Slide-Seq V2, files with count matrices of the appropriate formats need to be generated and imported into Scanpy data format by the user.

Let $G = \{g_{f,g}\}$, denote an $F \times G$ matrix of reference cell type signatures, which consist of $f = \{1,..,F\}$ gene expression profiles $G_{f,:}$ for $g = \{1,..,G\}$ genes, representing average expression of each gene in each cell type in linear mRNA counts space (not log-space). This matrix needs to be provided to cell2location and can be estimated from scRNA-seq profiles. Two strategies how to estimate this matrix are presented in Section 2.

Cell2location models the elements of $D$ as Negative Binomial distributed, given an unobserved expression level (rate) $\mu_{s,g}$ and a gene-specific over-dispersion parameter $\alpha_g$ which accounts for variance in expression of individual genes that is not explained by the reference cell type signatures:

$$d_{s,g} \sim \texttt{NB}(\mu_{s,g}, \alpha_g). \tag{1}$$

This is equivalent to a Poisson likelihood (measurement model) with a Gamma-distributed mean (expression model)[2]. Here NB distribution is parameterised by a Poisson rate parameter $\mu$ and over-dispersion $\alpha$, which corresponds to shape parameter of Gamma distribution in

Gamma-Poisson mixture

$$d_{s,g} \sim \texttt{Poisson}(\texttt{Gamma}(\alpha_g, \alpha_g/\mu_{s,g})). \tag{2}$$

The expression of level of genes $\mu_{s,g}$ in the mRNA count space is modelled as a linear function of expression signatures of reference cell types:

$$\mu_{s,g} = \underbrace{m_g}_{\text{technology sensitivity}} \cdot \underbrace{\left(\sum_f w_{s,f}\, g_{f,g}\right)}_{\text{cell type contributions}} + \underbrace{l_s + s_g}_{\text{additive shift}} . \tag{3}$$

Here, $w_{s,f}$ denotes regression weight of each reference signature $f$ at location $s$, which can be interpreted as the number of cells per location $s$ expressing each each reference signature $f$; $m_g$ denotes a gene-specific scaling parameter, which adjusts for global differences in expression estimates between technologies; $l_s$ and $s_g$ are gene- and location-specific components that capture additive shift which can be interpreted as contaminating RNA such as free-floating RNA.

The prior distributions on the unobserved parameters are as follows:

1. **Absolute abundance of cell types across locations.** The regression weights $w_{s,f}$ for each reference signature are modelled using a hierarchical Gamma prior

$$w_{s,f} \sim \texttt{Gamma}(mu = \mu^w_{s,f}, \sigma^2 = \mu^w_{s,f}/v^w), \tag{4}$$

where the prior mean $\mu^w_{s,f}$ accounts for linear co-variance / dependency between locations of different cell types and $v^w$ controls the prior strength. The mean parameter $\mu^w_{s,f}$ is modelled in a hierarchical fashion, decomposing the regression weights into $R$ latent groups of cell types $r = \{1, .., R\}$, thereby accounting for linear dependencies in spatial abundance between reference signatures:

$$\mu^w_{s,f} = \sum_r z_{s,r}\, x_{r,f}. \tag{5}$$

The number of latent groups $R$ is specified by the user (default $R = 50$). Intuitively, these $R$ groups of cell types can be thought of as cellular compartments in the tissue, which are characterized by shared cell type abundance profiles. We observed that the sensitivity of mapping low abundance cell types in particular increases when accounting for these dependencies (see Fig S2).

While the scaling parameter $m_g$ facilitates the integration across technologies, it leads to non-identifiability between $m_g$ and $w_{s,f}$, unless the informative priors on both variables are used. To address this, the prior distributions of $z_{s,r}$ and $x_{r,f}$ are defined to control absolute scale of the cell type abundance estimates $w_{s,f}$, guiding $w_{s,f}$ to the scale of the number of cells per location $s$ expressing each cell type reference signature $f$. In addition, these prior distributions help control the sparsity of how many cell types $f$ are expected at each location $s$, facilitating application of cell2location to tissues and technologies with varying numbers of cells and cells types per location. The hyper-parameters controlling the $w_{s,f}$ prior can be estimated from the paired histology image and the literature about the tissue (Section 1.3), whereas technology sensitivity $m_g$ can be hard to estimate *a-priori.*

- The total spatial abundance $z_{s,r}$ of latent cell type groups $r$ across locations $s$ is defined as Gamma distributed with a prior controlling the scale and sparsity of this distribution:

$$z_{s,r} \sim \texttt{Gamma}(Y_s/R, 1/(N_s/Y_s)), \tag{6}$$

  where $N_s$ corresponds to the average number of cells in each location, and $Y_s$ is the number of latent groups $r$ present in each location $s$ - two parameters that need to be specified by the user (See eq. (7)-(8) and section 1.3). The prior on $z_{s,r}$ is constructed such that $\sum_r z_{s,r}$ on average equals to the number of expected cells per location $\sum_r z_{s,r} = N_s$, and that on average each location has high $z_{s,r}$ for $Y_s$ expected cell type groups.
  Unobserved $Y_s$ and $N_s$ parameters are modelled as Gamma-distributed:

$$N_s \sim \texttt{Gamma}(mu = \hat{N}, \sigma^2 = \hat{N}/v^n) \tag{7}$$

$$Y_s \sim \texttt{Gamma}(mu = \hat{Y}, \sigma^2 = \hat{Y}), \tag{8}$$

  where $\hat{N}$ is a user-defined estimate of the expected number of cells per location; $\hat{Y}$ is the user-defined average number of cellular compartments / zones per location; and $v^n$ denotes the prior strength. Recommendation on setting there hyper-parameters are outlined in section 1.3.
- The latent variable $x_{r,f}$ represents the contribution of each latent cell type group $r$ to the abundance of cell types $f$ and is Gamma distributed with a prior controlling the number of cell types $f$ that have high values of $x_{r,f}$ in each group $r$:

$$x_{r,f} \sim \texttt{Gamma}(K_r/R, K_r), \tag{9}$$

  where $K_r$ represents the unobserved number of cell types for each group $r$. This prior controls the absolute values of $x_{r,f}$ such that on average $\sum_r x_{r,f} = 1$.
  $K_r$ is Gamma-distributed with a prior informed by user input:

$$K_r \sim \texttt{Gamma}(mu = \hat{A}/\hat{Y}, \sigma^2 = \hat{A}/\hat{Y}), \tag{10}$$

  where $\hat{A}$ is a user-provided average number of cell types per location; and $\hat{Y}$ is a user-provided average number of cellular compartments / zones per location (See recommendations in section 1.3). This prior tells that each group $r$ has large $x_{r,f}$ for many cell types $f$ when $\hat{A} > \hat{Y}$, and $\hat{A} = \hat{Y}$ indicates that the spatial abundance of each cell type $f$ is independent from other cell types.

2. **Gene-specific multiplicative scaling factor** $m_g$ is modelled as Gamma-distributed (Eq. (11) - (13)) with hierarchical prior $\alpha^m$ and $\beta^m$ reflecting the prior belief about the difference in sensitivity of single cell and spatial technologies

$$m_g \sim \texttt{Gamma}(\alpha^m, \beta^m) \tag{11}$$

$$\alpha^m \sim \texttt{Gamma}(mu = \mu_m^2/\sigma_m^2, \sigma^2 = \mu_m^2/\sigma_m^2) \tag{12}$$

$$\beta^m \sim \texttt{Gamma}(mu = \mu_m/\sigma_m^2, \sigma^2 = \mu_m/\sigma_m^2). \tag{13}$$

Here, $\mu_m$ denotes a user-provided average change in sensitivity, $\sigma_m^2$ denotes a user-provided deviation of the sensitivity of individual genes from $\mu_m$. The link between these hyper-parameters and priors $\alpha^m, \beta^m$ is set as described in the notation section 5.1. Section 1.3 details how these hyper-parameters are derived from data.

3. **Additive shift for genes and locations.** $l_s$ and $s_g$ are modelled as Gamma distributed with hierarchical priors on shape and rate of their distributions eq. (14) - (16):

$$l_s \sim \mathtt{Gamma}(\alpha^l, \beta^l) \tag{14}$$

$$s_g \sim \mathtt{Gamma}(\alpha^s, \beta^s) \tag{15}$$

$$\alpha^l, \beta^l, \alpha^s, \beta^s \sim \mathtt{Gamma}(1, 1) \tag{16}$$

4. **Overdispersion for each gene.** Containment prior[3] is used for modelling unobserved variance using NB over-dispersion $\alpha_g$. The idea of a containment prior is that, by the prior, NB distribution should be maximally close to Poisson distribution, which is achieved by putting most probability mass on large $\alpha_g$. So, under the following prior most genes have low over-dispersion:

$$\alpha_g = 1/o_g^2 \tag{17}$$

$$o_g \sim \mathtt{Exponential}(\beta^o) \tag{18}$$

$$\beta^o \sim \mathtt{Gamma}(mu, \sigma^2) \tag{19}$$

where constants $mu$ and $\sigma^2$ are provided by the user (See section 1.3).

In addition to reporting absolute cell density we also compute absolute number of mRNA molecules ($u_{s,f}$) contributed by each cell type $f$ to each location $s$:

$$u_{s,f} = w_{s,f}(\sum_g m_g \, g_{f,g}) \tag{20}$$

$u_{s,f}$ are a more robust estimate than either absolute cell density $w_{s,f}$ or relative cell proportions $w_{s,f}/\sum_f w_{s,f}$. For this reason it can be used as a criteria to accurately exclude non-mapped cell types (Fig S9). For comparison of cell and mRNA abundance in the mouse brain see Suppl Files 1 and 6.

#### 1.2 Multi-experiment extension of cell2location model

Joint modelling of multiple spatial data sets, where $e = \{1, .., E\}$ denote individual experiments, such as 10X Visium chips (i.e. square capture areas) and Slide-Seq V2 pucks (i.e. beads), provides the several benefits due to sharing information between experiments:

1. Increasing accuracy by improving the ability of the model to distinguish low sensitivity $m_g$ from zero cell abundance $w_{r,f}$, which is achieved by sharing the change in sensitivity between technologies $m_g$ across experiments. Similarly to common practice in single cell data analysis, this is equivalent to regressing out the effect of technology but not the effect of individual experiment.

2. Increasing sensitivity by sharing information on cell types with co-varying abundances via the variable $x_{r,f}$ which represents the latent groups $r$ of reference cell types $f$ with similar spatial abundance, and it's hierarchical prior, $N_r^x$ (the number of cell types per group).

The same reference cell type signatures $g_{f,g}$ are used for all experiments. Variables that serve as hierarchical priors are also shared between experiments (single value for each variable): $\alpha^m$, $\beta^m$, $\alpha^l$, $\beta^l$, $\alpha^s$, $\beta^s$, $\beta^o$.

When analyzing multiple spatial data sets jointly, the over-dispersion parameter $\alpha_{e,g}$ and additive background parameter $s_{e,g}$ are fit separately for each experiment as well as genes. Specifically, $\alpha_{e,g}$ depends on $o_{e,g}$ which is modelled as independent and exponentially distributed (eq. (21) - (22)) and $s_{e,g}$ is modelled as iid Gamma (eq. (23)):

$$d_{s,g} \sim \texttt{NB}(\mu_{s,g}, \alpha_{e,g} = 1/o_{e,g}^2) \tag{21}$$

$$o_{e,g} \sim \texttt{Exponential}(\beta^o) \tag{22}$$

$$s_{e,g} \sim \texttt{Gamma}(\alpha^s, \beta^s) \tag{23}$$

##### 1.3 Selecting hyper-parameters

Unless stated otherwise, we recommend the following strategy to select priors for every tissue and technology:

1. **Cell number $\hat{N}$ per location**, a single global estimate, is estimated from histology images paired to spatial expression data $d_{s,g}$ by examining the image using 10X Loupe browser (10X Visium data, Fig S6). We observed this number varies substantially between tissues (Fig S10). When the histology image is not available, the size of capture regions relative to the size of cells is used to estimate $\hat{N}$ (Slide-Seq V2).

   When the exact number of cells in each location can be obtained by using image segmentation to count nuclei (See Section 1.5), it can be provided as a location-specific $\hat{N}_s$ hyper-parameter and $v^n$ should be increased to make the prior more informative ($v^n = 10$). However, for all analysis in this manuscript a single global estimate was used. When the size of cell capture areas varies, such as in laser capture microscopy (LCM) method, we recommend to provide a location-specific $\hat{N}_s$ estimate.

2. **Number of cell types $\hat{A}$ per location**, a single global estimate, is chosen to be less than number of cells $\hat{N}$ and using background knowledge about the complexity of the tissue: spatial interlacing of tissue regions suggest high $\hat{A} \sim= 0.8\hat{N}$, locations dominated by a single cell type suggest $\hat{A} << \hat{N}$.

3. **Number of tissue zones $\hat{Y}$ per location**, a single global estimate, is chosen to be less than number of cell types $\hat{A}$ and is estimated using background knowledge about the complexity of the tissue: presence of spatially interlacing tissue zones (high complexity, $\hat{Y} \sim= 4$), discrete regions with dominant cell types (low complexity, $\hat{Y} = 1$). When location patterns of all cell type in the tissue are known to be spatially distinct $\hat{Y} = \hat{A}$ should be specified.

4. **Average difference in technology sensitivity $\mu_m$ and $\sigma_m^2$** parameters in eq. (12)-(13) are chosen by comparing average total number of mRNA per cell in the reference cell type data to the average total number of mRNA per location in the spatial data divided by $\hat{N}$. This process will be automated in future versions of cell2location.

The model will give more accurate absolute cell abundance estimates when these priors are chosen well, whereas the estimate of relative cell abundance and mRNA count from each cell type $u_{s,f}$ is quite robust to this choice. We observed 8 cells and 6 cell types per location in the mouse brain 10X Visium (Fig S6, Fig S10), but just 1-2 cells and cell types in the human brain and heart (not shown, using published spatial data). In the lymphoid and epithelial tissues, 10X Visium locations contain on average 30 cells and 10 cell types per location reflecting high spatial interlacing of cell type locations in these tissues (Fig S10 and unpublished).

Slide-Seq V2 locations are expected to have fractions of cells in each location ("bead"), where we observed 1-2 cell types per location (Fig S10)[4]. However, we observed large absolute cell abundance $w_{s,f}$ in the Slide-Seq V2 data (Fig 2H) potentially indicating that additional technical effects such as difference in sensitivity between "beads" need to be accounted for.

The following priors should be used at their default values unless stated otherwise:

1. **The number of co-located cell type groups is** $R = 50$. Regardless of selected $R$ the model tends to find much lower number of groups (3-7) $r$ with substantial values of $x_{r,f}$.

2. **Hyper-parameter** $v^w$, controlling the extent to which the co-abundance prior controls $w_{s,f}$, is set to a medium value of $v^w = 5$. Increasing $v^w$ forces stronger linear dependencies in $w_{s,f}$ leading to over-smoothing and lower model accuracy. Decreasing $v^w$ leads to cell types $f$ modelled as independent, which also decreases model accuracy because of the extent of those effects in real data.

3. $v^n$ **controlling the strength of** $\hat{N}$ **prior** in eq. (8) is set to 1 when a global rather than location-specific estimate $\hat{N}$ is specified.

4. **Hyper-parameters** $mu = 3$ **and** $sd = 1$ in eq. (19) are chosen to scale the exponential distribution in eq. (18) to put most probability density at low variance (large over-dispersion $\alpha_g$). The inference of this hyper-prior appears to be fairly robust to the choice of $mu$ and $sd$.

#### 1.4 Inference

Variational Bayesian Inference is used to approximate the posterior, building on the Automatic Differentiation Variational Inference (ADVI) framework implemented in pymc3[5]. Briefly, within this framework, the posterior distribution over unknown parameters are approximated by transformed (to ensure a positive scale) univatiate normal distributions. Inference is achieved by maximising log-likelihood of the data and minimising KL divergence from the posterior to prior, which are combined in the evidence lower bound (ELBO loss function). Training is stopped when ELBO stops increasing, usually after 20,000 - 40,000 iterations of training using ADAM optimiser (learning rate 0.005) with gradient clipping (total gradient norm $< 200$). Posterior mean, standard deviation, 5% and 95% quantiles for each parameter were computed using 1,000 samples from the Variational posterior distribution. We observed that 5% quantile provides a slightly more accurate estimate of cell abundance compared to the mean (data not shown).

Two or more restarts are performed to evaluate the stability of the inferred posterior distribution. Under default parameters we empirically observed that the inference results are very stable and hence model selection across restarts is not used by default and rather as a quality control step. In particular, significant divergence of the solutions between restarts tends to indicate poor match of the cell type reference or large mismatch between the prior distribution of $d_{s,g}$ (sample from the model given hyper-parameter choices) and the count distribution in the observed data $d_{s,g}$ (prior predictive check). Optionally, the suitability of the selected hyper-parameters can be assessed using prior predictive check (also provided within the cell2location package). As an additional control step, we recommend evaluation of the model fit as described in the tutorial: 1) by comparing the posterior mean estimate $log10(\mu_{s,g} + 1)$ to observed data $log10(d_{s,g} + 1)$ (posterior predictive check); 2) by evaluating consistency of inferred $w_{s,f}$ parameters between the training restarts.

##### 1.5 Nuclei Segmentation (optional)

To perform nuclei segmentation, H&E-stained image of the Visium-profiled tissue was used. The image was acquired using Hamamatsu slide scanner as described in the section on experimental processing. Full resolution images were divided into sub-images of size about 1000 by 1000 pixels. Then a state-of-the-art convolutional neural network (CNN)-based segmentation pipeline described in Caicedo et al.[6] was used. The model is available in GitHub repository selimsef/dsb2018_topcoders and the code to use it in yozhikoff/segmentation. An ensemble of 32 pre-trained CNNs each with Unet- or FPN-like architecture was used to classify pixels to 3 classes: background, nuclei and nuclei boundaries. Predicted segmentation masks were averaged across CNNs, then a watershed algorithm was applied to refine predicted boundaries and separate individual nuclei. As a final step, a gradient boosting model over morphological features such as nuclei colour, shape and size was used to remove false-positively detected nuclei. Obtained individual nuclei masks were used to compute several morphological features: 1) area (count of pixels in a mask), 2) shape - ratio of lengths of the major and minor axis of an ellipse fitted to the mask, and 3) position - center of mass of the image containing individual mask. Kd-tree approach was used to efficiently assign nuclei to Visium locations. Kd-tree implementation from the scikit-learn package was used. Nuclei positions in the histology image are reported in Suppl. Data.

#### 2 Estimation of reference expression signatures of cell types

We provide two methods for deriving reference expression signatures of cell types $g_{f,g}$ from scRNA-seq expression data $J = \{j_{c,g}\}$ of each gene $g = \{1..G\}$ in cell $c = \{1..C\}$. For both methods, we assume that cell type labels for each cell are provided by the user, for example as obtained using conventional cell clustering to identify putative cell types and sub-populations.

First, untransformed and unnormalised count matrix $J_{c,g}$ is filtered to select expressed genes (Fig S1) at 2 cut-offs: 1) selecting genes detected at mRNA count $> 0$ in many cells ($> 5\%$ of cells), 2) selecting genes detected at mRNA count $> 0$ in a few cells ($5\% >$ cell count $> 10$) but with large mean expression across non-zero cells ($> 1.1$). The cut-offs listed are a guideline, with values adjusted for every data set (Methods).

Next, we offer two alternative methods to derive $g_{f,g}$ from the scRNA-seq count matrix. Although method 1 is computationally more efficient, we recommend method 2 as robust general-purpose solution:

1. **Computing mean count each gene $g$ in cell cluster $f = \{1..F\}$.** This method is computationally highly efficient and generally yields good mapping quality as long as the scRNA-seq reference data are obtained from a single batch. We also recommend this methods when using Smart-Seq 2 scRNA-seq data, as the distributional assumptions of method 2 are not appropriate for this technology.

2. **Model-based estimation of reference expression signatures of cell types $g_{f,g}$ using a regularised Negative Binomial regression.** This model robustly derives reference expression signatures of cell types $g_{f,g}$ using the data composed of multiple batches $e = \{1..E\}$ and technologies $t = \{1..T\}$. Adapting the assumptions of a range of computational methods for scRNA-seq[7,8], we assume that the expression count matrix

$J = \{j_{c,g}\}$ follows a Negative Binomial distribution with unobserved expression levels (rates) $\mu_{c,g}$ and a gene-specific over-dispersion $\alpha_g$:

$$J_{c,g} \sim \mathtt{NB}(\mu_{c,g}, 1/\alpha_g^2). \tag{24}$$

We model $\mu_{c,g}$ as a linear function of reference cell type signatures and technical effects:

$$\mu_{c,g} = (g_{f,g} + b_{e,g})\, h_e\, p_{t,g}. \tag{25}$$

Here, $h_e$ denotes a multiplicative global scaling parameter between experiments/batches $e$ (e.g. differences in sequencing depth); $p_{t,g}$ accounts for multiplicative gene-specific difference in sensitivity between technologies; $b_{e,g}$ accounts for additive background shift of each gene in each experiment $e$ (proxy for free-floating RNA).

All model parameters are constrained to be positive. We use a weak L2 regularisation of $g_{f,g}$, $b_{e,g}$ and $\alpha_g$, and a strong penalty for deviations of $h_e$ and $p_{t,g}$ from 1. $g_{f,g}$ is initialised at analytical average for each cell type $f$. $b_{e,g}$ is initialised at average expression of each gene $g$ in each experiment $e$ divided by a factor of 10. Such informative initialisation leads to fast convergence.
Pytorch implementation of training using mini batches makes the model fast (2min-15min on GPU) and scalable to very large data sets ($>$ 100k cells). Maximum a posteriori optimisation is used to find parameters of this model with batch size of 1024 cells, ADAM optimiser learning rate 0.01. The number of epochs needed for training is determined using cross-validation performed on held-out 10 percent of cells of each cell type $f$.

To verify that the model successfully accounted to non-biological sample effects $h_e$, $p_{t,g}$ and $b_{e,g}$, a corrected expression matrix $J_{c,g}^{corrected}$ is obtained by normalising $J_{c,g}$:

$$J_{c,g}^{corrected} = J_{c,g}/(h_e\, p_{t,g}) - b_{e,g} \tag{26}$$

$J_{c,g}^{corrected}$ matrix is subjected to standard Scanpy workflow[1] to verify the mixing of the cells from different samples and technologies within matching cells types (Fig S17B).

### 3 Comparison of cell2location with alternative methods

Cell2location is compared to alternative methods, which have recently been proposed or developed in parallel (Table 1). The methods are compared based on the modelling approach (points 1-2), the method for estimating expression signatures of cell types (3), whether the methods account for key features of spatial (3-4) and scRNA-seq reference data (4-6), support joint modelling of multiple spatial experiments / batches (7), whether the methods provide features relevant for scalability such as mini-batch training and GPU acceleration (8-9), programming language (10).

### 4 Downstream analyses

The output of cell2location can feed into a range of downstream analysis approaches. Here, we outline strategies of how to combine cell2location with cell clustering, as well as non-negative matrix decomposition to identify groups of co-located cell types.

Table 1: Comparison of cell2location with other alternative methods

| Criteria | cell2location | stereoscope[9] | Seurat V3[10] | SpotLIGHT[11] | NNLS (autogenes)[12] | RCTD[4] |
|---|---|---|---|---|---|---|
| 1. Data distribution, normalisation | Negative Binomial, non-negative parameters | Negative Binomial, non-negative parameters | Normalised log(x+1) | Counts, non-negative parameters | Counts, non-negative parameters | Negative Binomial, log-normal model |
| 2. Inference method | Variational Inference | Penalised maximum likelihood | PCA + MNN | NNLS | NNLS | Penalised maximum likelihood |
| 3. Method for estimating signatures | Method provided but user-derived signatures can be used (See Section 2) | Hard-coded method (NB regression with unusual parametrisation) | Not needed | Hard-coded method (topic model) | User-derived signatures | Hard-coded method (mean gene expression) |
| 4. Accounting for the difference in sensitivity between single cell and spatial technology | Yes | Yes | Unclear | No | No | Yes |
| 5. Models similarity of locations between cell types | Yes | No | No | No | No | No |
| 6. Method in #3 accounts for technology and batch effects in scRNA data | Yes | No | Yes | No | No | No |
| 7. Joint modelling of multiple spatial experiments | Yes | No | Unclear | No | No | No |
| 8. GPU acceleration | Yes | Yes | No | No | No | No |
| 9. Mini-batch training | At lower accuracy | Yes, likely at lower accuracy | No | No | No | No |
| 10. Language | Python | Python | R | R | Autogenes in Python, other in R | R |

#### 4.1 Identifying tissue regions by clustering

The cell abundance $w_{s,f}$ estimated by cell2location model is used to describe similarity of locations $s$ by constructing a KNN graph that represents connectivity of locations in terms of their cell composition. Standard implementation of Leiden clustering (function scanpy.tl.leiden) in the Scanpy package[1] can utilise cell composition KNN graph to group organisation of the tissue. Leiden clustering was performed with default arguments except for resolution (see main methods section). Obtained regions were cross-referenced with the mouse brain anatomy using corresponding histology images and similar clusters were merged to generate the broad region map.

#### 4.2 Groups of co-located cell types

To identify cellular compartments of the tissue by utilising cell abundance estimates $w_{s,f}$, we applied the non-negative matrix factorisation model (NMF). Absolute cell abundance $w_{s,f}$ of each cell type $f$ across locations $s$ is modelled as an additive function of the cell type groups $r$ (cellular compartments). This model is similar to the co-abundance prior in the main cell2location model (eq. (4)), however we observed that when applied as a downstream analysis step, the factorisation tended to identify more granular cellular compartments. Under additive decomposition, several cellular compartments can be present in one location providing a major practical and conceptual advantage over discrete clustering (Fig S16B). Cell abundance estimates $w_{s,f}$ are modelled as an additive decomposition:

$$w_{s,f} = \sum_{r} z_{s,r}\, x_{r,f} \tag{27}$$

where $x_{r,f}$ represents NMF weight describing the contribution of each cell type group $r$ to cell types $f$; $z_{s,r}$ represents NMF weight describing the abundances of each cell type group $r$ across locations $s$.

Similarly to cell2location model (eq. (20)), we compute $u_{s,r}$ which can be intuitively interpreted as the number of cells from each cell type group $r$ in each location $s$ and use it for visualisation:

$$u_{s,r} = z_{s,r} \sum_{r} x_{r,f} \tag{28}$$

Inference is performed using standard NMF in the scikit-learn package[13]. The model is trained 5 times to evaluate stability of the identified cellular compartments. The first training iteration is always selected.

**Rationale for applying the NMF to multi-cell spatial data.** We hypothesise that linear dependencies in the abundance of cell types across locations reflects the interaction between these cell types, specifically, any interaction that could affect mutual location of these cell types. For this reason we tested this approach on human lymph nodes, a tissue where locations of cell types are determined by their interactions (Fig 4D). 10X Visium technology combines cell types located within the $55\mu m$ diameter of capture areas, a distance at which the action of local paracrine signals and direct adhesive contacts can be observed, likely driving linear dependencies in cell abundance. We assume that the signal does not propagate between neighbouring locations spaced by $45\mu m$ gaps, therefore using proximity information is not essential. Therefore we note that that this interpretation of NMF decomposition of cell abundances is a unique feature of grid-based technologies that aggregate multiple cells and cell types within single locations. This interpretation does not directly apply to technologies with increased resolution (Slide-Seq V2) or technologies where capture regions represent selected features.

Potential cell interactions driving linear dependencies in cell type abundance can be classified into:

- Signals promoting proliferation and survival of other cell type
- Adhesion molecules that increase the frequency of contacts
- Recruitment signals such as CXCR5 / CXCL13[14]
- Repulsive signals, inhibiting the action of chemo-taxis signals such as PD-1 / PD-L1 interactions[15]
- Interactions that induce new transcriptional states

### 5 Appendix

#### 5.1 Notation

Two alternative parameterizations of Gamma distribution are used: 1) $\alpha$ (also called shape or over-dispersion) and $\beta$ (also called rate); 2) $\mu$ and $\sigma^2$; with the link between the two given by:

$$\mu = \alpha/\beta; \sigma^2 = \alpha/\beta^2 \tag{29}$$

$$\alpha = \mu^2/\sigma^2; \beta = \mu/\sigma^2 \tag{30}$$

By default, $\alpha$ and $\beta$ parametrization is used; when $\mu$ and $\sigma^2$ are used their notation is shown.
